# Target Dependent Coordinated Biogenesis Ensures Cascaded Expression of miRNAs in Activated Murine Macrophage

**DOI:** 10.1101/2021.06.11.448041

**Authors:** Susanta Chatterjee, Ishita Mukherjee, Mainak Bose, Shreya Bhattacharjee, Saikat Chakrabarti, Suvendra N. Bhattacharyya

## Abstract

MicroRNAs (miRNAs) repress protein expression by binding to the 3’ UTR of the target mRNAs. By exploring the effect of target mRNA on biogenesis of cognate miRNAs, we have noted miRNA with higher number of binding sites (primary miRNA) coordinates the biogenesis and activity of another miRNA (secondary miRNA) having binding sites on the 3’ UTR of a common target mRNA. From the quantitative data obtained from macrophage cells, we detected miR-146a-5p as a “primary” miRNA that coordinates biogenesis of “secondary” miR-125b, miR-21 or miR-142-3p to target new sets of mRNAs to balance the immune response in activated macrophage cells. Interestingly, target dependent coordinated biogenesis of miRNAs, happening on the rough endoplasmic reticulum attached membrane, ensures a cumulative mode of action of primary and secondary miRNAs on the secondary target mRNAs where a cascaded effect of primary miRNA on its secondary targets has been detected. Extensive computational analysis for the presence of coordinated biogenesis pairs of miRNAs in mammalian cells has also allowed us to construct a coordinate biogenesis repository to determine context specific coordinated biogenesis relationships exists for specific pairs of miRNAs in mammalian cells.

**Graphical Abstract:** 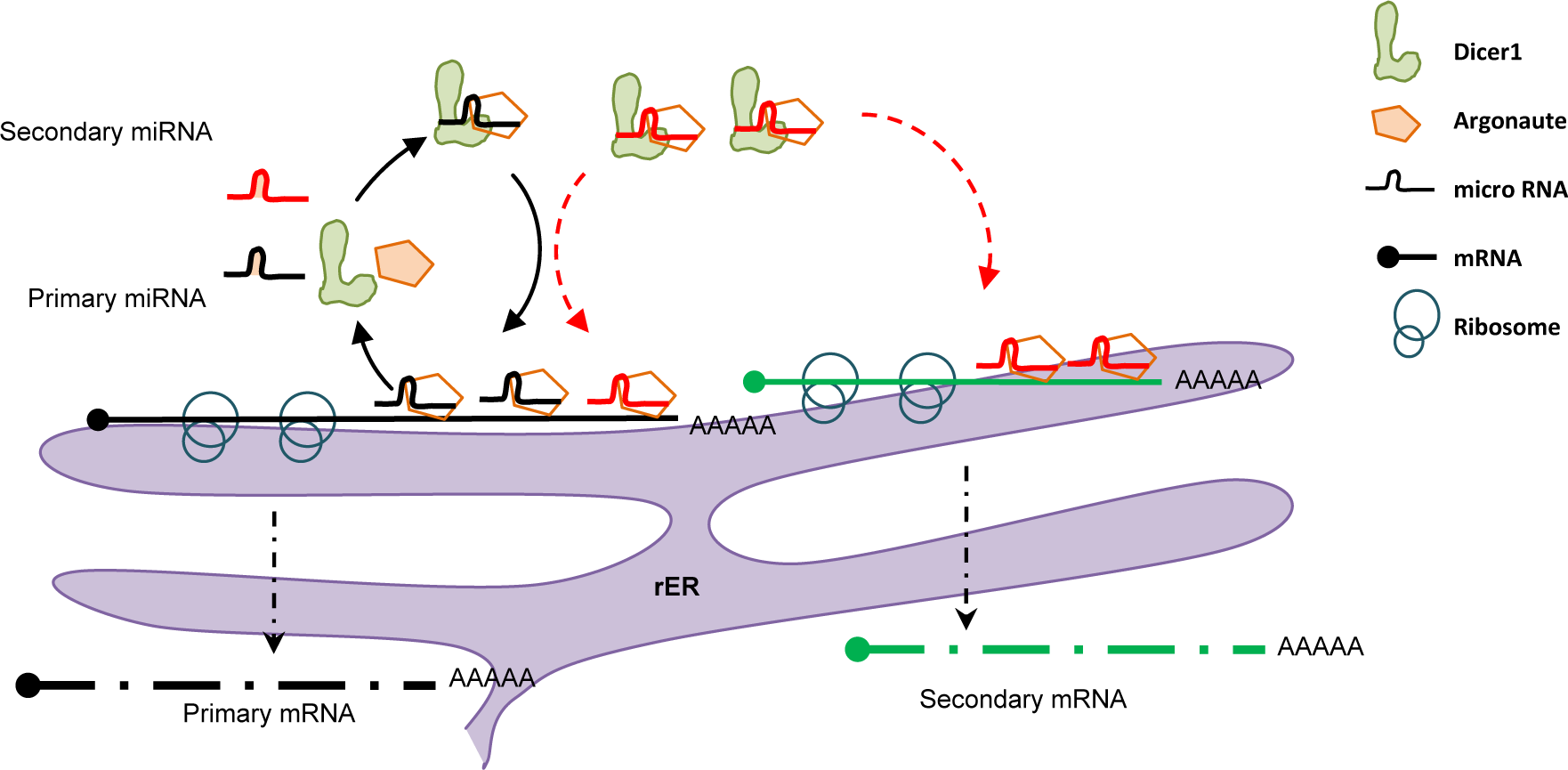

- miRNA with higher number of binding sites (primary miRNA) can coordinates the biogenesis and activity of another miRNA (secondary miRNA)
- Target dependent coordinated biogenesis of miRNAs ensures a cumulative mode of action of primary and secondary miRNAs on the secondary target mRNAs on rER attached polysomes
- miR-146a-5p acts as a “primary” miRNA to coordinate biogenesis of “secondary” miR-125b, miR-21 or miR-142-3p in activated macrophage cells
- Coordinate biogenesis balance the immune response in activated macrophage cells by ensuring propagation of primary miRNAs effect to diverse target mRNAs through secondary miRNAs.

## Introduction

MicroRNAs (miRNAs)are small regulatory RNAs that are primarily processed from introns, exons or intergenic regions of the mammalian genome by the canonical or non-canonical pathway of miRNA biogenesis (O’Brien et al., 2018). After sequential processing, the precursor form of the miRNAis cleaved by the endonuclease Dicer1 to form miRNA/miRNA* duplex and the miRNA encoding strand gets loaded onto Argonaute proteins to form RNA induced silencing complex (miRISC) for silencing of target mRNAs (Bartel, 2004). Conventionally, miRNA binds to the target mRNA with 5’ conserved seed sequence complementarity and represses the expression of proteins by translational repression and/or destabilization of target mRNA.

Intracellular levels of different miRNAs produced during development in specific cell types or in response to specific environmental cues may be governed by transcription factors, post-transcriptional regulatory mechanisms influencing miRNA processing, or abundance of precursor mRNA (Bose and Bhattacharyya, 2016; O’Connell et al., 2010). In a reciprocal fashion, miRNA activity and abundance can be altered by multiple factors including RNAs which themselves are under miRNA regulation. Group of coding and non-coding RNAs [pseudo genes, long noncoding RNAs (lncRNA), circular RNA (circRNA)] can modulate the miRNA activity by forming an integrative network (Chan and Tay, 2018). This interconnected regulatory crosstalk gives rise to the idea of competitive endogenous RNA(ceRNA) hypothesis where ceRNA with miRNA binding sites (also referred as miRNA responsive elements, MRE) competes for miRNA binding against the endogenous target mRNAs that ultimately leads to derepression of the target mRNA through “miRNA sponge effect” of ceRNA (Salmena et al., 2011). Apart from the well-established network of this mode of regulation of PTEN mRNA, a critical tumour suppressor encoding mRNA regulated by various group of ceRNAs (Poliseno et al., 2010; Tay et al., 2011), the functional sequestration of miR-122 in hepatic cells by a Hepatitis C Virus (HCV) synthetic construct also manifest the “sponge-effect” to elicit functional de-repression of miR-122 targetome (Luna et al., 2015). Although ceRNA mediated derepression could be exacerbated by overexpression of different synthetic RNA transcripts, the concept is debatable due to lack of abundant physiological levels of ceRNAs in cellular context (Denzler et al., 2016).

There is ample evidence for different modes of regulatory control of RNA over miRNA biogenesis and activity. Kleaveland et al. have shown a non-coding RNA (ncRNA) network where different species of ncRNA cooperatively act to modulate neuronal activity in the brain (Kleaveland et al., 2018). It is also reported how increase in number of the binding sites on a target mRNA increases the ‘processivity’ of Dicer1, leading to enhanced cognate miRNA biogenesis from precursor per unit time in hepatocytes subjected to amino acid starvation and subsequent re-feeding (Bose et al., 2017). We were interested to dissect the consequent effect of this regulation by target mRNA of a specific miRNA on other miRNAs with *cis*-binding sites on the 3’ UTR of the same target mRNA. Is there existence of a cooperative biogenesis of miRNAs that share binding sites on 3’ UTR of common target mRNAs? Or in contrast, if there exists a competitive binding among groups of miRNAs on 3’ UTR of common target mRNAs?

To unravel the mystery, we wanted to explore a physiologically relevant system where multiple miRNAs may have a crosstalk to fine-tune common signalling pathways. TLR4-activated inflammation, elicited by exogenous or endogenous ligands, does give rise to several acute and chronic diseases, and plays a critical role as an amplifier of the inflammatory response. Expression profiling in human monocytes revealed groups of endotoxin-responsive miRNAs which fine tune the expression of signalling mediators during inflammatory escalation (O’Neill et al., 2011; Taganov et al., 2006). miRNAs can stimulate or regulate various signalling pathways such as TLR-signalling pathways (TLR4, TLR3, and TLR7/8 etc.), NF-kβ pathway or MAPK pathways (MAPK/ERK, MAPK/JNK, and MAPK/p38 pathways) to mount pro- and anti-inflammatory responses or modulate innate immune responses (Momen-Heravi and Bala, 2018; Nejad et al., 2018).

We anticipated the existence of a molecular coordination among those miRNAs teaming up to regulate the immune response. We hypothesize that the coordination might be achieved by target mRNAs affecting the biogenesis of multiple miRNAs to create a regulatory module in infected macrophages.

We validated our hypothesis of coordinated miRNA biogenesis in mammalian cells using a reporter model, where miRNA let-7a could influence the biogenesis and activity of exogenously expressed miR-122 in a target mRNA-dependent miRNA biogenesis process. Further, by adopting the endotoxin-activated macrophage system, we documented the cooperative biogenesis of miRNAs. We have observed that miR-146a-5p could modulate biogenesis and activities of groups of miRNAs like, miR-125b, miR-142-3p and miR-21. In turn, this phenomenon resulted in repression of secondary target mRNAs that are controlled by the newly generated secondary miRNAs. These network-based regulations of miRNAs allowed us to propose a Target Dependent Cooperative Biogenesis (TDCB) in mammalian cells where miRNA species with higher number of binding sites influences the biogenesis and activity of a group of “cooperative” miRNAs having target sites on the same common mRNAs. The process is driven by a primary mRNA-miRNA driven compartmentalization of secondary miRNAs and their targets to the rough endoplasmic reticulum (rER) membrane associated polysomes and required for TDCB. Considering this observation, a comprehensive computational analysis has also been performed to suggest a general prevalence of the coordinated biogenesis (CB) phenomenon for multiple miRNAs both in human and mouse model. Our observation also suggests, this “epistatic” mode of regulation of miRNA could in turn influence and fine tune different cellular signalling pathways for re-imposition of cellular homeostasis.

## Results

### Stepwise expression of miRNAs in activated macrophage cells

Bacterial membrane lipopolysaccharide (LPS) is an endotoxin synthesized by gram-negative bacteria that serves as an immediate activator of TLR4 signalling pathway to increase the production of proinflammatory cytokines which protect host cells by innate immune response (Lu et al., 2008). In activated macrophage cells, the miRNA activity loss due to miRNA uncoupling from Ago proteins precedes the de *novo* miRNA biogenesis and re-establishment of the miRNA mediated repression of target cytokine mRNAs during prolonged exposure to LPS (Goswami et al., 2020; Mazumder et al., 2013). In this process, re-association of Ago2 with housekeeping miRNAs was ensured (i.e. let-7a miRNA) while increased expression of several new miRNAs was also noted (O’Neill et al., 2011; Zhou et al., 2010). Interestingly, majority of these miRNAs were induced in a phase wise manner and the transcriptional surge of the precursors encoding these miRNAs has not been sufficient to explain the cascaded expression pattern observed for these miRNAs in mammalian macrophage cells. A group of miRNAs were proven to be the key mediators of TLR4-responsive pathways and some of them like miR-21 and miR-146a-5phad showed dose dependent increase after LPS-induced activation of murine macrophage cells (Figure 1A)(Quinn and O’Neill, 2011).These miRNAs also showed a specific temporal expression pattern after LPS exposure. While miR-155 shows a peak of expression at an early time-point, miR-146a-5p and miR-21 get induced at comparably later time-points(Figure 1B) (Kurowska-Stolarska et al., 2011; Sheedy et al., 2010; Taganov et al., 2006). Increase of specific miRNAs in LPS-treated primary macrophage or PEC (peritoneal exudate cells) treated with LPS was also documented (Figure 1C). The induced expression of these miRNAs is associated with reduced expression of their targets in macrophage after 24h of LPS exposure confirming the functionally active forms of the induced miRNAs (Figure 1D).

**Figure 1.**
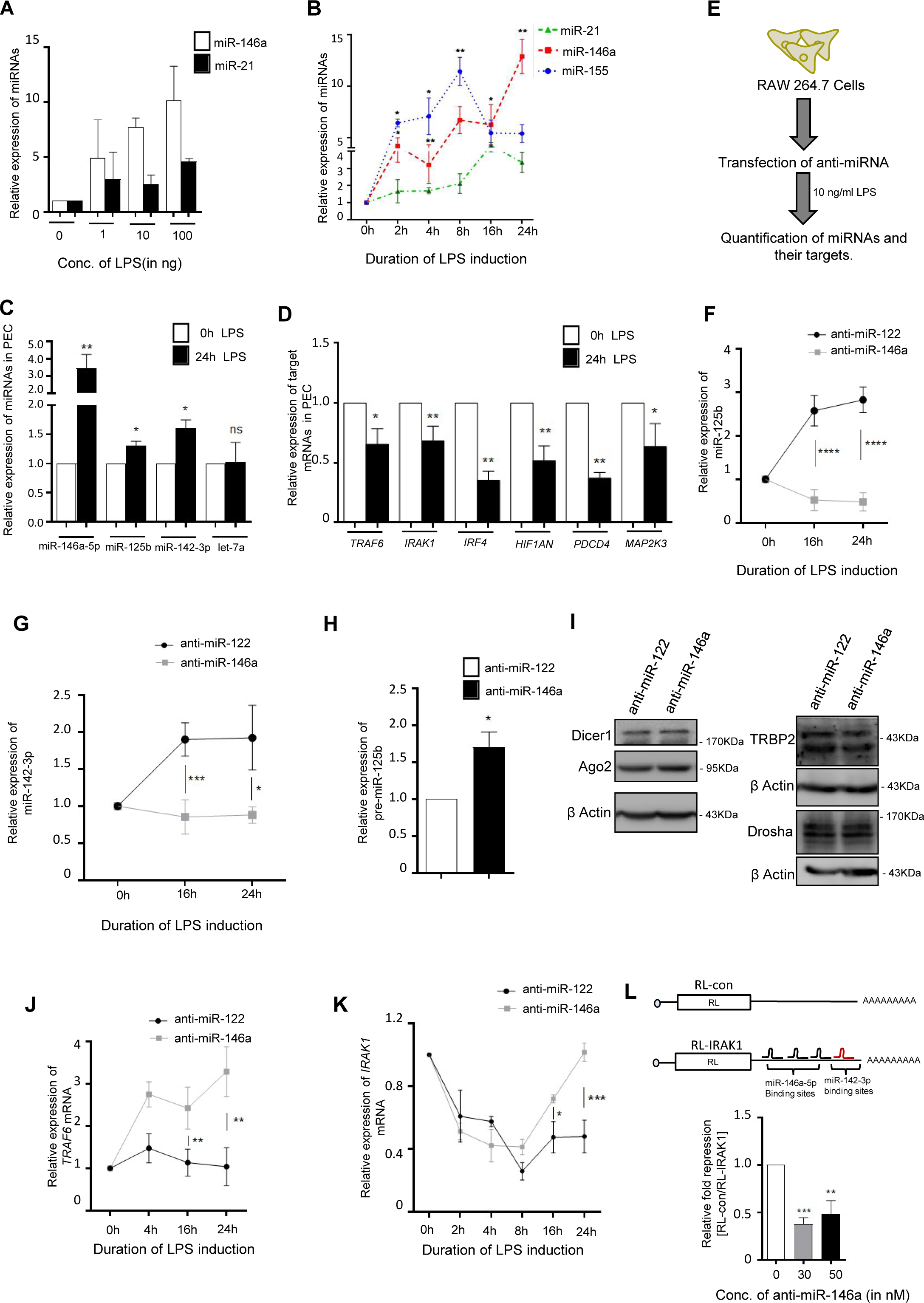
miR-146a regulates biogenesis of specific secondary miRNAs in LPS-induced murine macrophages. **A** Expression of miR-146a-5p and miR-21 in murine macrophages after their treatment with increasing concentration of LPS. PCR data reveals dose-dependent increase of miR-146a-5p and miR-21 expression level after LPS treatment for 24h in RAW 264.7 cells (n=3). **B** Changes in the levels of different miRNAs against time during LPS (10ng/ml) treatment. qRT-PCR data is showing an increase of miR-155, miR-146a-5p and miR-21 expression after LPS induction. Levels of miR-155 reached a peak of expression at 8h post-infection whereas levels of miR-146a-5p and miR-21levels increased further post 8h time point (n=3). **C-D** LPS treatment of PEC cells increase the expression of LPS-responsive (miR-146a-5p, miR-125b, miR-142-3p) but not the non-responsive (let-7a) miRNAs. Treatment was done with 10ng/ml of LPS for 24 h (C). Along with this, levels of respective targets of those miRNAs; TRAF6 & IRAK1 (of miR-146a), IRF4 & HIF1AN (of miR-125b) and PDCD4 & MAP2K3 (of miR-21) were measured and plotted after 24h of LPS treatment. The data obtained thus confirms the enhanced activities of those miRNAs that are unregulated with LPS treatment (D) (n=4). **E** Schematic representation of experiments. RAW 264.7 murine macrophage cells were either transfected with 30 nM anti-miR-146a or anti-miR-122 (control) oligos. Post-transfection, cells were induced with 10ng/ml LPS for respective time-points (n=3). **F-G** miR-125b (F) and miR-142-3p (G) levels after LPS induction in anti-miR-146a or anti-miR-122 transfected RAW 264.7 murine macrophage cells. qRT-PCR data shows both the miRNAs were significantly downregulated post 16h & 24h of LPS treatment after anti-miR-146a treatment compared to anti-miR-122 control (n=3). **H** Cellular level of pre-miR-125b in anti-miR-146a or anti-miR-122 transfected RAW264.7 cells treated with LPS for 24h (n=3). **I** Cellular levels of miRNP-associated proteins in anti-miR-146a or anti-miR-122 transfected LPS treated macrophages. miRNP-associated proteins & processing enzymes Dicer1, Ago2, TRBP2 & Drosha were western blotted to detect no change in their expression in miR-146a-5p inactivated LPS treated RAW246.7 cells. **J-K** Expression ofTRAF6 (J) & IRAK1 (K) mRNA in anti-miR-146a and anti-miR-122 transfected macrophage cells after 10ng/ml of LPS treatment over 0-24h time (n=3). **L** Schematic representation of RL reporter mRNA with 3’UTR of IRAK1 mRNA with respective miRNAs binding sites is shown in upper panel. Renilla luciferase reporter assay for RL-IRAK1 reporter in anti-miR-146a or anti-miR-122 transfected LPS treated macrophage cells. Renilla reporters were co-transfected with firefly expression plasmids and firefly expression was used as transfection control (n=3). 18s rRNA or GAPDH and U6 snRNA level has been used as endogenous target for Real time qRT-PCR mRNA and miRNA quantification respectively. Student’s t-tests were used for all comparisons. p < 0.05 (*); p < 0.01 (**); p < 0.001(***); p < 0.0001 (****). Values are means from at least three biological replicates ± SEM.

### miR-146a-5p mediates biogenesis of groups of secondary miRNAs

Along with miR-146a-5p, expressions of miR-125b and miR-142-3p were also upregulated after LPS induction (Figure 1 E-G)(Murphy et al., 2010; Sun et al., 2011; Zhou et al., 2010). Interestingly, we could observe a significant decrease of miR-125b and miR-142-3p level in LPS treated macrophages after transfection with anti-miR-146a (Figure 1 F-G). In contrast, pre-miR-125b showed increased accumulation correlating with decreased processing of miR-125b from its precursor in anti-miR-146a treated cells (Figure 1H). We also observed reduced miR-21 levels even after LPS induction in anti-miR-146a treated macrophages while miR-21 levels remained unchanged after miR-155 inhibition, used as a negative control (Figure S1A). Let-7a expression, taken as a non-cooperative miRNA control, remained unaltered post-LPS induction (Mazumder et al., 2013) and level of let-7a did not change even after miR-146a inhibition (Figure S1B). To understand the molecular players behind this differential miRNA expression, we wanted to investigate expression levels of the different miRNA associated protein factors viz. Dicer1, Ago2 and TRBP that are known to be associated with processing of pri- and pre-miRNAs. We could not observe any change in expression of miRNP-formation associated protein factors, negating the possibilities of miR-146a-5p mediated upregulation of pan-miRNP-processing event to alter generic miRNA levels in activated macrophages (Figure 1L). These observations reflect the specific regulatory role of miR-146a, as a primary miRNA, on biogenesis of specific secondary miRNAs such as miR-142-3p, miR-125b or miR-21, in LPS stimulated macrophage cells.

### miR-146a-5p mediates biogenesis of secondary miRNAs requires mRNA targets having bindings sites for both primary and secondary miRNAs

miR-146a-5p is one of key miRNAs that are induced in macrophages after LPS exposure. As discussed earlier, miR-146a-5p is a critical immune regulator that acts as a molecular brake on inflammation and malignant transformation (Magilnick et al., 2017). miR-146a-5p regulates Toll-like receptor and cytokine signalling via a negative feedback regulation loop mediated through down-regulation of IL-1 receptor-associated kinases 1 (IRAK1) and TNF receptor-associated factor 6 (TRAF6) protein levels (Figure 1J-K) (Taganov et al., 2006). Multiple TLR4 signalling components TRAF6 or IRAK1 encoding mRNAs bearing multiple miR-146a-5p binding sites, showed synchronized expression with miR-146a-5p level in a time-dependent manner and showed derepression after miR-146a-5p specific inhibition in LPS-induced macrophage cells (Figure 1J-K) (Hou et al., 2009). The irrefutable function of miR-146a-5p on combating inflammation and orchestrated expression of multiple miR-146a-5p target mRNAs led us curious to explore the effect of miR-146-5p on miRNAs that shares the 3‘ UTR target sites with miR-146a-5p and are known as important regulators of innate immune response also. Target dependent biogenesis of miRNAs has been reported in mammalian cells (Bose and Bhattacharyya, 2016). The 3’UTR of most of the endogenous target mRNAs have binding sites for several miRNAs and among them some miRNAs may have multiple binding sites for a single target mRNA. Is it possible that miRNAs with higher number and strong binding sites can positively regulate the binding and target driven biogenesis of “secondary” miRNAs that are otherwise expressed low in macrophage cells?

Different predicted (TARGETSCAN) and experimentally validated (miRTARBASE) databases and previous publication based combinatorial analysis of TLR4 pathway-miRNA network study revealed that TRAF6 bears one conserved miR-125b binding site and IRAK1 possesses one conserved miR-142-3p binding site along with previously mentioned miR-146a-5p binding sites which is present on both targets. This is conserved across mammals in the phylogenetic tree (Figure 1L, S1C-D) (Agarwal et al., 2015; Sun et al., 2011; Wang et al., 2014).Inhibition of miR-146-5p in macrophage cells derepresses the miR-142-3p mediated repression of RL reporter mRNA with 3’UTR of *IRAK1.* Confirming the loss of miR-142-3pactivity in absence of active miR-146-5p to repress the IRAK1 (Figure 1L).miR-21 is another critical modulator of LPS-responsive pathways (Sheedy et al., 2010).TARGETSCAN and previous reports also reveal 3’ UTR of both the TRAF6 and IRAK1 mouse mRNA contains single miR-21-5p binding sites along with miR-146a-5p (Figure S1C-D) (Agarwal et al., 2015; Nara et al., 2019).

### Target driven coordinated biogenesis of secondary miRNAs is coupled with cooperative repression of their targets in murine macrophage

To understand the differential repressive activities of probable “cooperative groups of secondary miRNAs” in primary miRNA-compromised cells, we introduced a plasmid encoding renilla luciferase (RL) reporter mRNA that contains MAD1L1-3′UTR with miR-125b binding site (Bhattacharjya et al., 2013). Luciferase assay showed reduced repression of miR-125b reporter target post-LPS induction in the anti-miR-146a transfected murine macrophage cells (Figure 2A). Ago2 is one of the key effector proteins of the RISC complex (Chendrimada et al., 2005). Ago2-based immunoprecipitation study showed significant reduction in association of candidate miRNAs viz. miR-125b and miR-142-3p as well as secondary target mRNAs of miR-125b (HIF1AN) with Ago2 in LPS-induced macrophages transfected with miR-146a inhibitor oligos (Figure 2B-C). miR-21 showed a similar trend of reduced Ago2 association in anti-miR-146a treated cells (Figure S1E).

**Figure 2.**
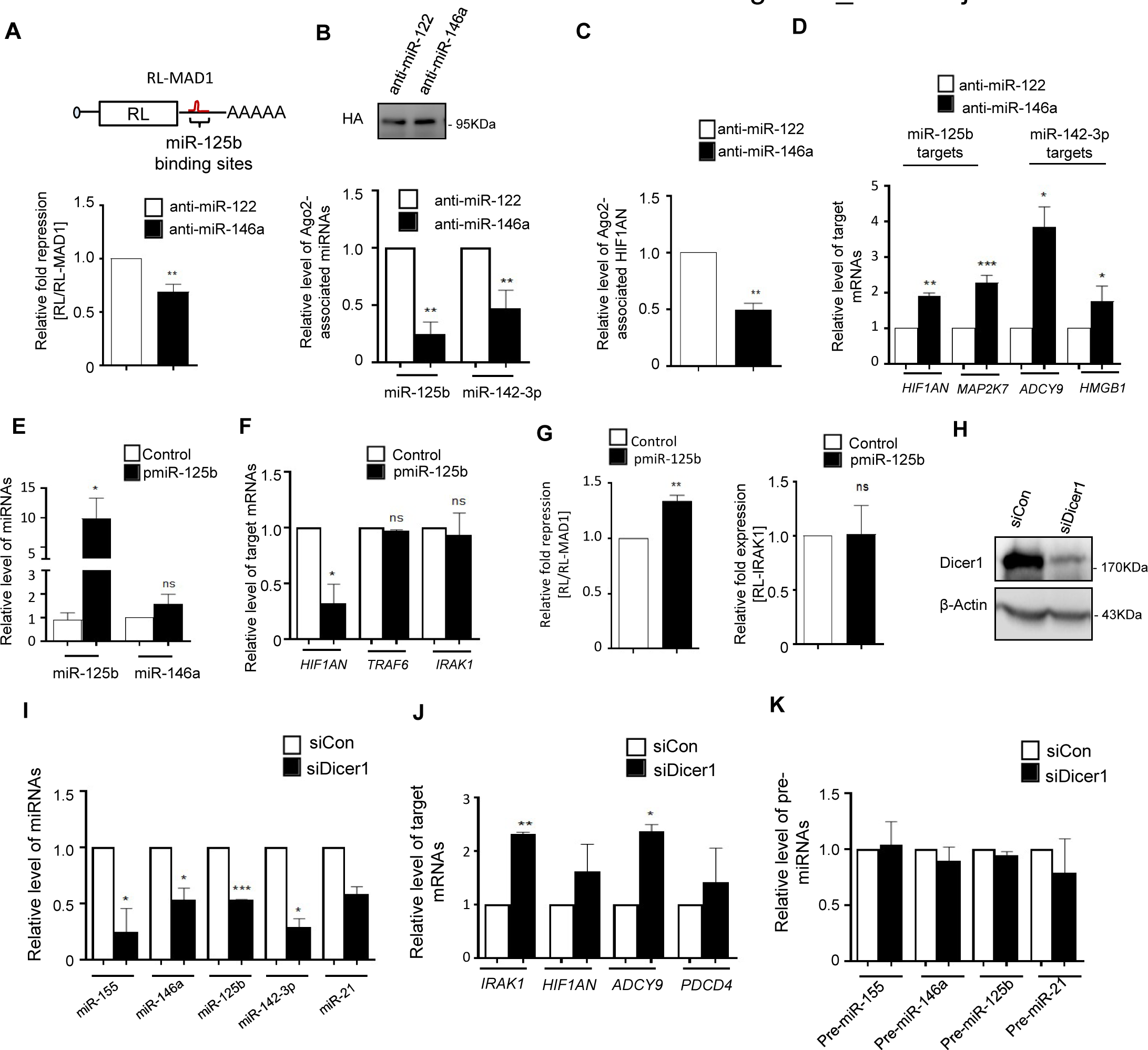
Coordinated biogenesis of secondary miRNAs driven by miR-146a-5p is linked with increased activity of secondary miRNAs in activated macrophage. **A** Dual luciferase assay shows reduction in miR-125b activity in anti-miR-146a transfected RAW 264.7 cells treated with LPS. Schematic representation of the RL-MAD1 luciferase mRNA that harbours one miR-125b binding sites on its 3’ UTR acts as reporter for miR-125b. A renilla reporter without any miRNA binding sites (RL-con) was used as a control. Firefly Luciferase (FF) construct without miRNA sites was used to normalize the transfection efficiency between the sets used for this assay (n=3). **B-C** Reduced association of miR-125b and miR-142-3p with Ago2 after 24h of LPS treatment of anti-miR-146a transfected cells(B). Reduced association of miR-125b target mRNA HIF1AN with Ago2 has also been observed(C). Macrophages were activated with LPS after transfection with HA-Ago2 expression construct and respective anti-miR inhibitor oligos. LPS induction was done for 24 h. Western blots of HA has confirmed the amount of Ago2 pull down post-immunoprecipitation that was used for normalization (n=3). **D**Derepression of miR-125b & miR-142-3p endogenous targets after 24h of LPS induction in anti-miR-146a transfected cells. qRT-PCR analysis of endogenous miR-125b target (HIF1AN &MAP2K7) and miR-142-3p target (ADCY9 & HMGB1) were done to estimate the relative mRNA content after anti-miR-146a transfection (n=3). **E** miR-125b does not affect cellular miR-146a-5p content. qRT-PCR data shows no effect of overexpression of miR-125b on miR-146a level in murine macrophage cells transfected with miR-125b expression plasmid pmiR-125b against pCIneo control plasmid transfected cells. **F-G** Unaltered activities of miR-146a-5p on target mRNAs in pmiR-125b transfected cells. Real-Time PCR data shows no significant change in thelevels of endogenous targets of miR-146a-5p, TRAF6 & IRAK1 after miR-125b overexpression. Decreased HIF1AN level confirmed increased activities of miR-125b after its overexpression (F). Dual luciferase data exhibits no detectable change in miR-146a reporter RL-IRAK1 expression but increased repression of miR-125b reporter RL-MAD1 after miR-125b overexpression from pmiR-125 plasmid. Firefly Luciferase (FF) constructs was used to normalize the transfection levels between the sets used for this assay (G) (n=3). **H-K** Knockdown of Dicer1 inhibits LPS induced upregulation of groups of LPS-responsive miRNAs. Western blots show confirmation of Dicer1 knockdown in LPS-treated murine macrophages (H). Western Blot shows reduced level of Dicer1 after siDicer mediated knockdown in RAW cells. β-Actin serves as endogenous control. qRT-PCR data shows reduced level miR-155, miR-146a, miR-125b, miR-142-3p and miR-21 in Dicer-knocked down cells treated for 4h with LPS (I). qRT-PCR data were used to detect the levels of respective precursor miRNAs after Dicer1-knock down in LPS-treated cells (J). Levels of target mRNAs of the respective miRNAs also showed derepression upon Dicer1 depletion by siDicer1 in LPS-activated macrophages (K). In all cases n=3. 18s rRNA or GAPDH and U6 snRNA level has been used as endogenous control for Real time qRT-PCR based mRNA and miRNA quantification respectively. Concentrations and duration of LPS induction on macrophages were 10ng/ml for 24h wherever not mentioned. Paired two-tailed Student’s t-tests were used for all comparisons. p < 0.05 (*); p < 0.01 (**); p < 0.001(***). In (A-G, I-K) values are means from three biological replicates ± SEM.

To explore the resonating effect of this cooperative phenomenon, we were interested to measure the effect on endogenous validated targets of those miRNAs generated as “secondary miRNAs” by the primary miRNA-target mRNA cooperative pairs and to quantify their expression. MAP2K7 and HIF1AN & ADCY9 and HMGB1 are experimentally validated targets for miR-125b & miR-142-3p respectively (Chen et al., 2014; Huang et al., 2009; Wang et al., 2016; Zhang et al., 2014). There were no experimentally validated or predicted miR-146a-5p binding sites on the 3’ UTRs of those mRNAs and thus their expression should not get affected directly by miR-146a-5p binding to the respective 3’ UTRs (Agarwal et al., 2015). Our observation revealed derepression of those “secondary” non-miR-146a-5p target genes upon miR-146a-5pinhibition in LPS treated macrophages (Figure 2D). Consistent with CB effect, two important targets of miR-21 viz. PDCD4 and MAP2K3 were also derepressed (Figure S1F) (He et al., 2019; Sheedy et al., 2010). Overall, the observations depict a probable non-canonical function of miR-146a-5p, where a primary miRNA i.e. miR-146a-5p could cooperatively modulate the levels of secondary miRNAs as well as their target mRNAs that does not harbour any miR-146a-5p binding site. This is an auxiliary pathway through which the miRNAs can affect the expression of their secondary miRNAs mediated by a target mRNA that has binding sites for both the primary and secondary miRNAs.

Does miR-146a-5p-mediated biogenesis of secondary miRNAs and coordinated repression of secondary mRNAs have a feedback response on “primary” miR-146a-5p level? We wanted to decipher if there exists a reciprocal effect of secondary miRNA miR-125b on miR-146a-5p mediated coordinated biogenesis function. To understand this, we introduced pmiR-125b in murine macrophages that expresses miR-125b exogenously and investigated the alteration of miR-146a-5p expression and activity. Dual luciferase assay confirms the increased miR-125b biogenesis and activity after miR-125b expression (Figure 2E-G). We could not detect any significant change in miR-146a-5plevel or its’ differential activity on respective endogenous targets or reporter mRNA after miR-125b expression. Thus, nullifying the probable miR-125b-mediated feedback responses on miR-146a-5p biogenesis or activity (Figure 2E-G). These findings led us to conclude thatmiR-146a-5p governed CB operates unidirectionally for miR-125b in activated macrophages. The network is specific to miR-146a-5pand differential expression of miR-125b-5pcould not lead to consequent changes of activity and biogenesis of the miR-146a-5p. miR-146a-5p mediated cooperativity is thus specific for a group of “secondary” miRNAs and it probably acts unidirectionally.

The RNAseIII endonuclease Dicer1 plays a prominent role in processing of mature miRNA from its precursor and it has been reported important for the target driven biogenesis of the miRNAs in mammalian cells (Bose and Bhattacharyya, 2016). To identify the role of Dicer1 in processing and generation of the LPS induced “secondary” miRNAs, we knocked down Dicer1 and did not observe LPS-induced induction of sets of secondary miRNAs in Dicer1 depleted cells (Figure 2H-I). This data is consistent with the derepression of respective target mRNAs in Dicer1 depleted cells (Figure 2J). However, the levels of precursor miRNAs do not change significantly for either primary or secondary miRNAs in Dicer1 depleted cells (Figure 2K). Thus, Dicer1 is required for induced expression of respective miRNAs that are expressed upon LPS treatment of macrophages.

### Target mRNA driven cooperative biogenesis of miRNAs also occurs in non-macrophage cells

To reconfirm the specificity and authenticity of our observation with miR-146a-5p and its downstream secondary miRNAs, we used let-7a and miR-122 as primary and secondary miRNA pair in HEK293 cells to test our hypothesis in non-immune mammalian cells. We designed a RL encoding reporter mRNAs bearing three bulged let-7a sites and a single miR-122 binding site while a RL mRNA with mutated let-7a sites but with a single miR-122 site to serve as control (Figure 3A). Let-7a is endogenously expressed in HEK293 cells and miR-122 expression was induced by doxycycline from a miR-122 expression plasmid with a doxycycline responsive promoter (Figure 3B)(Bose and Bhattacharyya, 2016). *RL-3xbulge-let-7a-1xbulge-miR-122*, but not the *RL-3xbulge-let-7amut-1xbulge-miR-122*, should be responsive to translation repression by let-7a, but repression by miR-122 should ideally be operative for both constructs. Interestingly, when miR-122 was induced, repression by miR-122 was noted prominently for the *RL-3xbulge-let7-1xbulge-miR-122* mRNA but not for the defective let-7a site-bearing construct (Figure 3C). Quite predictably, the let-7a sites on *RL3xbulge-let7-1xbulge-miR-122* induced higher biogenesis of mature let-7a compared to the one with mutated recognition sites (Figure 3D). Interestingly, miR-122 biogenesis was greatly enhanced by *RL3xbulge-let7-1xbulge-miR-122*mRNA but not by the mutated let-7a binding sites containing construct. For further interception of the process and to confirm let-7a binding dependent biogenesis of functional miR-122 miRNPs via a common reporter target mRNA, miR-122 sites have been cloned downstream of a RL encoding sequence while the primary GFP mRNA having binding sites for both let-7a (3x bulge) and miR-122 (1x bulge) was used to induce target-driven miR-122 expression in HEK293 cells (Figure 3E). Repression of miR-122 RL reporter mRNA suggests decreased expression of *RL* mRNAs with bulged or perfect miR-122sites in *GFP-3xBulge-let-7a -1xbulge miR-122* mRNA co-expressing cells (Figure 3F-G). Increased miR-122 production by *GFP-3xbulge-let-7a-1xbulge-miR-122* was further evident from the reduction in total mRNA levels of endogenous let-7a targets (*K-Ras, N-Ras*) as well as miR-122 targets (*CAT-1, ALDO A, and GYS1*) in cells expressing constructs with both let-7a binding and miR-122 binding sites(Figure 3H). Thus, TDCB of miRNAs is a widespread phenomenon and not limited to macrophage cells exclusively. Along with, this also suggests a cooperativity of action amongst miRNAs to coordinate the biogenesis and activity of miR-122 at secondary level by primary let-7a miRNA that is mediated by their shared target mRNAs forming a regulatory effect in mammalian cells.

**Figure 3.**
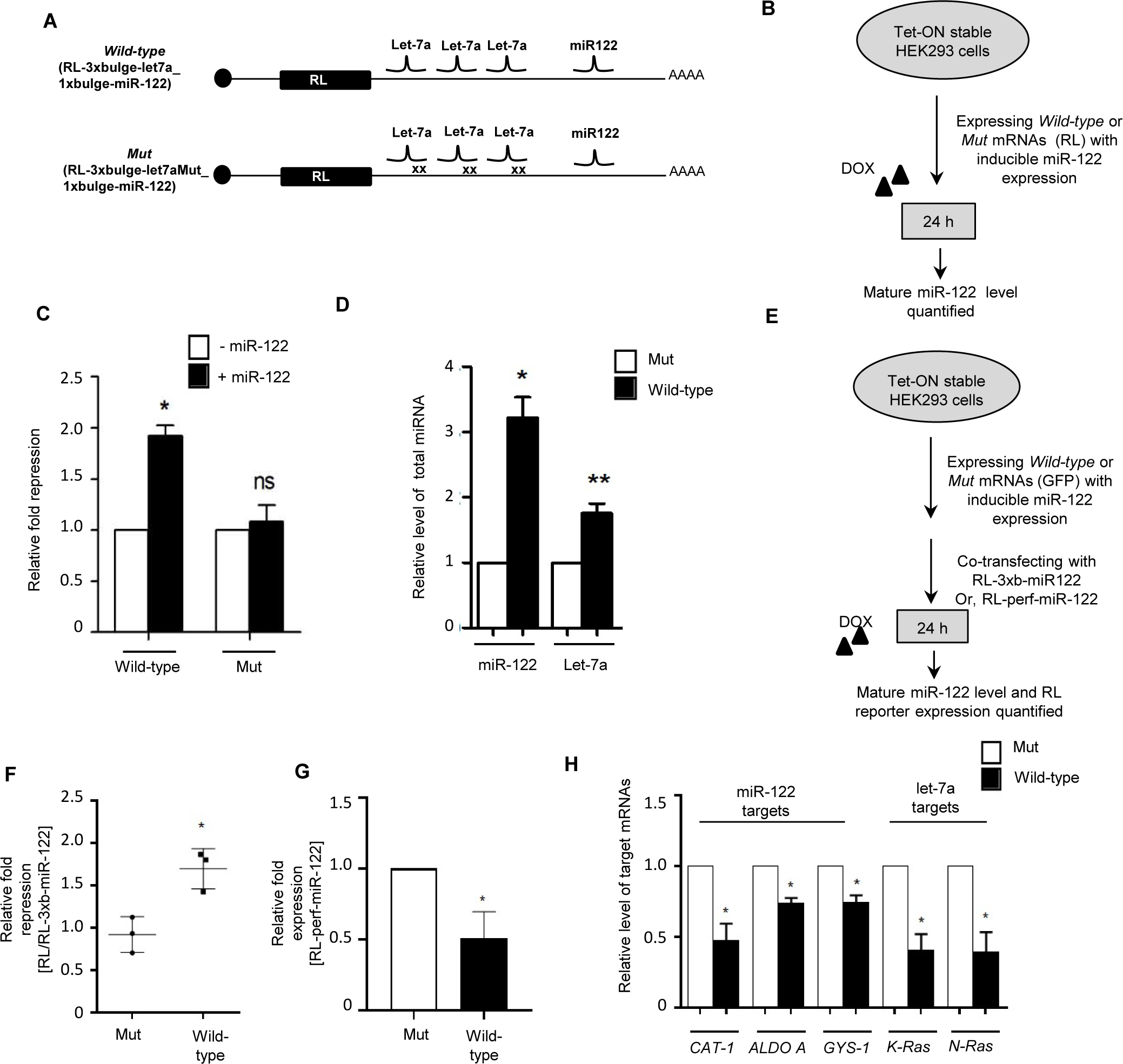
Target driven coordinated biogenesis leads to co-regulation of gene expression in non-macrophage cells. **A** Design of the reporter constructs used for testing coordinated biogenesis of miRNA by its targets in non-macrophage HEK293 cells. RL reporters with wild type or mutant let-7a miRNA binding sites and miR-122 sites in their 3’ UTRs are shown. **B** Scheme of the experiments to test coordinated biogenesis of miRNAs in HEK293 cells. **C** miR-122 represses target mRNA bearing functional let-7a sites but not with mutated let-7a sites present in cis with a single miR-122 site. HEK293 cells were transfected with RL-3xbulge-let7a_1xbulge-miR-122 or RL-3xbulgemut_1xbulge-miR-122, with or without pre-miR-122, and fold repression measured by luciferase assay done with above mentioned reporters and a renilla reporter without any miRNA binding sites (RL-con). Relative fold repression values have been plotted taking the values without miR-122 expression as unit in both cases. **D** Target dependent increased biogenesis of let-7a and miR-122 after transfection with let-7a target site containing RL-3xbulge-let7a_1xbulge-miR-122 construct but not with the let-7a mutant sites bearing construct in HEK cells expressing pre-miR-122. U6 snRNA was used as endogenous control (n=3). **E-G** Experimental scheme to study effect of coordinated biogenesis of secondary miR-122 by GFP reporter mRNAs with both let-7a and miR-122 sites on expression level of miR-122 Renilla reporter mRNAs (E). Dual luciferase reporter assay shows increased repression and decreased expression of secondary reporter gene bearing only miR-122 binding sites in presence of GFP reporter constructs containing intact let-7a binding sites compared to the mutated one with defective let-7a binding sites for RL reporters with imperfect miR-122 (F) or perfect miR-122 (G) sites (n=3). **H** Target driven miRNA biogenesis operates for both let-7 and miR-122 miRNAs simultaneously in presence of common target mRNA as evident from repression of their respective endogenous targets. The miR-122 targeted *CAT-1, ALDO A* and *GYS-1* and let-7a targets *K-Ras* and *N-Ras* showed repression in cells transfected with reporter mRNAs with both let-7a and miR-122 sites. (n=3). 18s rRNA or GAPDH and U6 snRNA level has been used as endogenous controls for Real time qRT-PCR based mRNA and miRNA quantification respectively. Paired two-tailed Student’s t tests were used for all comparisons. p < 0.05 (*); p < 0.01 (**); p < 0.001(***). In (B-D, F-H) values are means from at least three biological replicates ± SEM. Pre-miR-122 inductions in HEK293 T cells were ensured by treatment of the cells with doxycycline for 24h.

### Target mRNA compartmentalization to rER ensure miRNA compartmentalization and CB biogenesis of secondary miRNAs

To understand the mechanistic detail of how the target mRNA with binding sites of both primary and secondary miRNAs may have a role to play in CB of secondary miRNAs, we have done a cell fractionation experiment to follow the subcellular localization of secondary miRNA and their targets in primary miRNA expressing cells. From our previous findings, we do know rER attached polysomes as the sites of miRNA biogenesis (Bose et al., 2020) and how the mRNA targeting to rER attached polysome precedes the miRNP interaction (Barman and Bhattacharyya, 2015). The repressed messages then subsequently get relocalized to endosomes for their degradation and uncoupling from miRNPs (Bose et al., 2017). We have done the fractionation experiment by separating the digitonin soluble cytoplasmic fraction from the insoluble rER enriched membranes (Bose et al., 2020) and have identified the enhanced accumulation of Renilla reporter mRNAs having wild type let-7a sites over the reporter with mutant sites when both of them harbour the one secondary miR-122 binding sites (Figure 4A-B). The polysome association of the target mRNA also enhanced for wild type let-7a sites containing reporter mRNAs (Figure 4B). Interestingly the secondary miRNA target (in this case GFP-3xB-miR-122 for secondary miR-122) and both the primary (let-7a) and secondary (miR-122) miRNAs also showed enhanced association with rER fraction in cells co-expressing RL reporter mRNA with wild types let-7a and miR-122 sites compared to let-7a mutant site containing reporter (Figure 4C-D). These results were consistent with enhanced primary let-7a and secondary miR-122 endogenous target mRNA association with rER in cells expressing *RL-3xbulge-let-7a-1xbulge-miR-122*, but not the *RL-3xbulge-let-7amut-1xbulge-miR-122*(Figure 4E). This data points to the importance of primary mRNA for the enhanced rER compartmentalization of both secondary miRNA and their targets expressed in a heterologous context in HEK293 cells.

**Figure 4.**
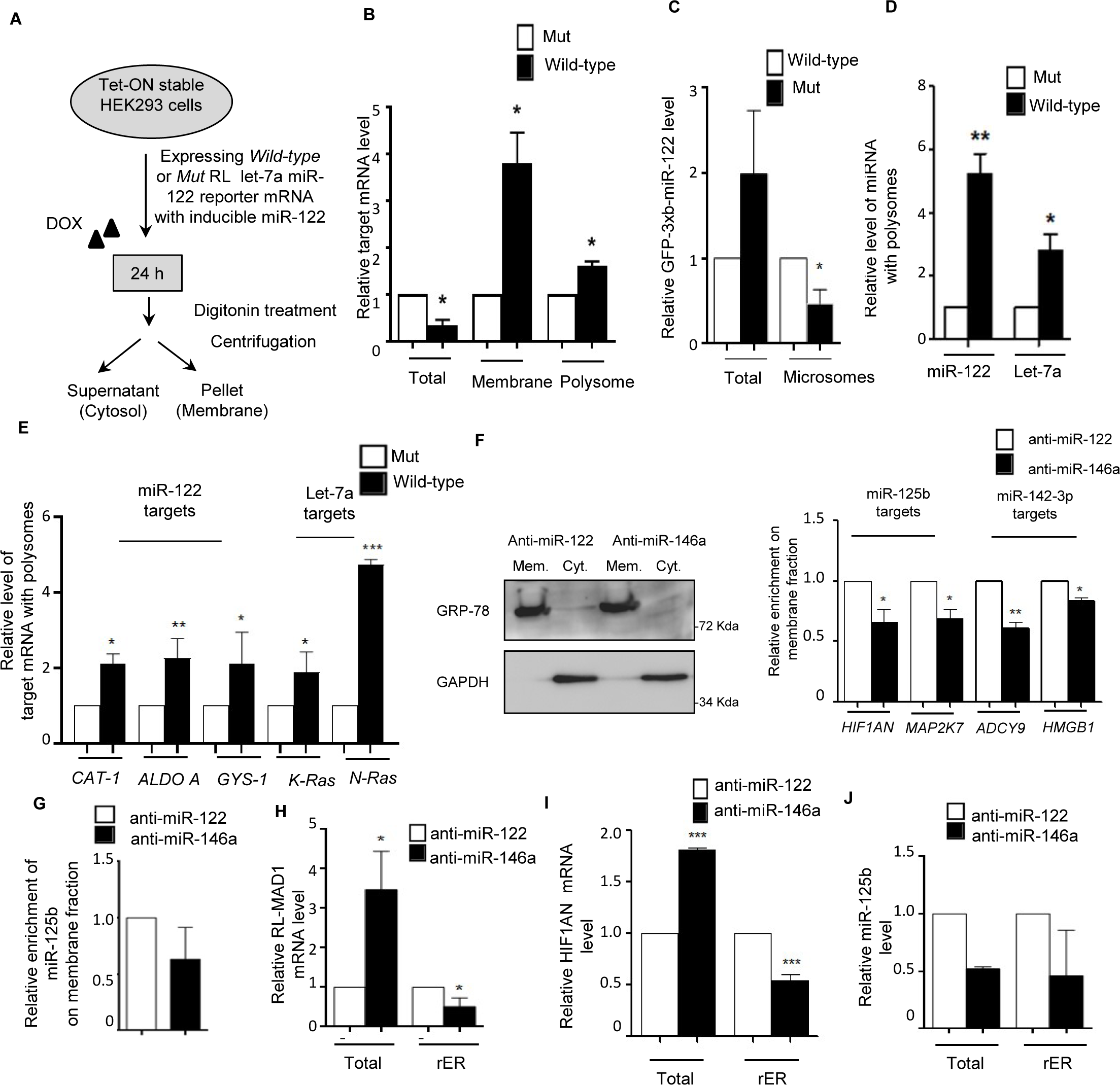
Target mRNA driven increased rER targeting of primary miRNAs drives increased biogenesis of CB miRNAs to repress its’ targets at rER membrane. **A** Schematic representation of the experimental set up. **B** Wild-type mRNA is more membrane associated and polysomal compared to the Mut version. miR-122 expression was induced by doxycycline for 24h in Tet-ON stable HEK293 cells transfected with target mRNA constructs and mRNA levels measured in total cell lysate, digitonin-insoluble membrane and polysomes. Values for Mut have been set as 1. **C** Increased cellular and decreased microsomal sequestration of secondary GFP reporter target mRNA for miR-122. **D-E** Target induced increased miRNAs and their endogenous targets are polysome enriched. Polysomes isolated have been used for quantification of mature miRNAs and target mRNAs. **F** Western Blot data of membranes and cytosolic marker proteins after digitonin based cellular fractionation. Western Blots of GRP78 and GAPDH acts as marker protein for membrane and cytosolic compartments respectively confirms proper isolation of respective compartments on anti-miR inhibitor transfected macrophages (left panel). Level of target mRNAs of respective secondary miRNAs (i.e. miR-125b & miR-142-3p) shows reduced membrane association on anti-miR-146a transfected LPS induced murine macrophages. **G** Digitonin mediated cell permeabilization study shows reduced membrane enrichment of secondary miRNA miR-125b on anti-miR-146a transfected LPS induced murine macrophages. **H-I** Reduced microsomal association of target mRNAs of miR125b on anti-miR-146a transfected LPS induced murine macrophages. Real-time PCR of endogenous target of miR-125b(HIF1AN) and reporter target of miR-125b (RL-MAD1L1) shows reduced microsomal sequestration along with increased level of respective derepressed mRNAs on cellular fractions. **J** Reduced microsomal enrichment of miR-125b on anti-miR-146a transfected LPS induced RAW264.7 cells. RT-PCR data shows miRNA level on microsomes also decreases as well as on total cellular fraction after anti-miR-146a transfection. 18s rRNA or GAPDH and U6 snRNA level has been used as endogenous target for Real time qRT-PCR mRNA and miRNA quantification respectively. Student’s t tests were used for all comparisons. p < 0.05 (*); p < 0.01 (**); p < 0.001(***). Values are means from at least three biological replicates ± SEM.

Do the membrane compartmentalization of secondary miRNA and their targets dependent on functionality of primary miRNA in RAW264.7 cells? We have tried the cell fraction assay with anti-miR-146 transfected RAW264.7 cells treated with LPS. We documented reduced membrane attachment of secondary miRNA, miR-125b, and its targets in anti miR-146a transfected RAW264.7 cells (Figure 4F-G). We have also observed a reduced rER attachment of a secondary RL reporter *RL-MAD1* mRNA, endogenous target mRNA *HIF1AN* and miR-125b itself with the rER in cells transfected with anti-miR-146 (Figure 4H-J).. This data suggest that the primary miRNA-driven CB is associated with compartmentalization of secondary miRNAs and their targets with the rER in mammalian cells and important for target repression.

### Coordinate biogenesis relationship within miRNA-mRNA interactome in lipopolysaccharide exposed macrophages

To explore the miR-146a-5p dependent and target mediated cooperativity in secondary miRNA biogenesis in a larger cellular context, we analysed the differential expression of whole cell transcriptome after LPS mediated stimulation of macrophages. We have observed that miR-146a-5p may exhibit CB wherein it influences the generation of a cluster of miRNAs. Based on certain probable pre-requisite conditions for CB phenomenon the possible CB regulator-target relationships of the CB-regulator mmu-miR-146a-5p in lipopolysaccharide exposed murine macrophages were predicted (Figure 5A). Based on the assumptions considered for the CB phenomenon to occur, around 65,535 CB regulator-target (miRNA-1: mRNA-A coupled to miRNA-2: mRNA-B) relationships were sampled. These CB regulator-target relationships were obtained across 57 genes (Gene A) and 187 of miRNA-2 considering mmu-miR-146a-5p (miRNA-1) as the CB-regulator. Utilizing expression information to ascertain which genes are likely to be expressed during macrophage immune responses, we have predicted the mmu-miR-146a-5p coordinate biogenesis regulatory network. Around 116 CB regulator-target relationships were determined in which both ‘Gene A’ and ‘Gene B’ were down-regulated given that the CB regulator and secondary effector miRNA in turn are likely to be up-regulated. However, 91 CB regulator-target relationships were considered for further analysis since there was no known or published direct regulatory relationship between ‘Gene A and B’ in those sets (Figure 5B). These criteria resulted in a final set of 91 predicted CB regulator-target relationships for possible experimental validation where both ‘Gene A’ & ‘Gene B’ are significantly down-regulated (Table S1). Thus, based on the assumptions of co-ordinate biogenesis with the help of our computational analysis, we have identified 21 possible candidate secondary effector miRNAs (i.e., miRNA-2) whose expression may be regulated by miR-146a-5p as a result of CB. Further, it is possible that these 21 secondary effector miRNAs may subsequently regulate the expression of 42 mRNA species (from secondary effector genes) (Table S1). In order to confirm our hypothesis, subsequently we have determined whether a change in the levels of the CB-regulator is reflected in the levels of the secondary effector miRNA and its corresponding target mRNA in biochemical experiments. In order to validate whether the proposed computational methodology can indeed identify biologically relevant CB regulatory network, candidate miRNAs were selected. Since Bach2 (as Gene A) possesses the highest number of miR-146a-5p binding-sites (4 in number), we have selected the associated set of CB regulator-target relationships for further analysis and validation. Our biochemical data shows significant down-regulation of two potential miRNAs (namely miR-16, miR-21-3p) after anti-miR-146a transfection in LPS induced macrophage cells (Figure 5C). Consequently, we could observe significant derepression in the predicted secondary effector genes as well, such as Mcm5, Ncapg2(downstream targets of miR-16) and Rrm2, Aspm, Ncadp3, Fen1 (downstream targets of miR-21-3p) (Figure 5D). Therefore, it is likely that a perturbation at the levels of the CB-regulator (miR-146a-5p) affects the down-stream targets in the coordinate biogenesis regulatory network, such as the secondary effector miRNA (miR-16, miR-21-3p and miR-142-3p as previously shown) and their target genes. This data suggests that miR-146a-5p can indeed potentially exhibit CB for a group of miRNAs and regulate their activity in LPS-induced murine macrophages.

**Figure 5.**
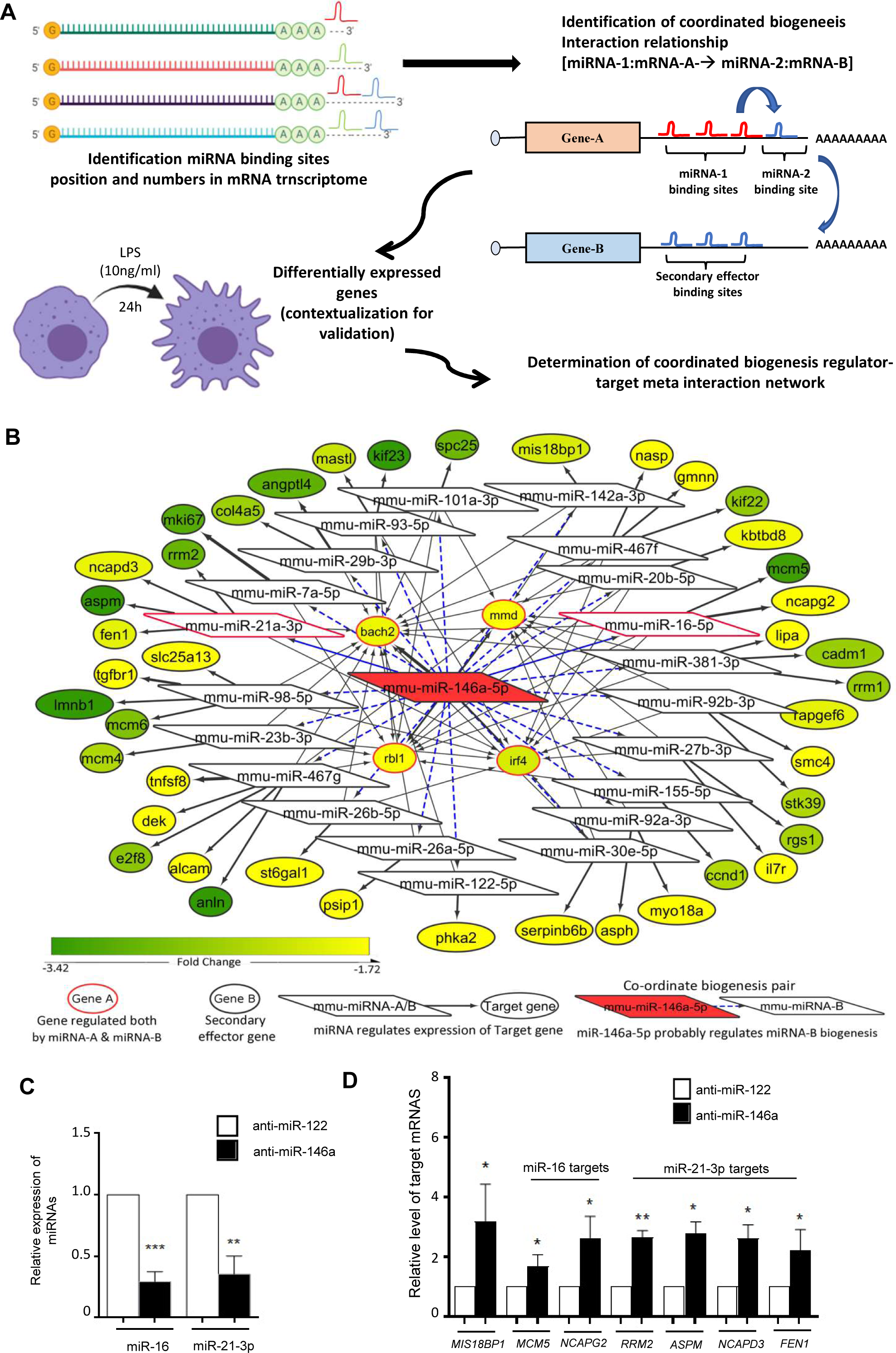
Network analyses and expression mapping reveals the group of miRNAs and their targets that are co-ordinately regulated by miR-146a-5p. **A** Schematic representation of the methodology utilized to predict CB regulator-target relationships for miR-146a-5p based on miRNA-mRNA network analysis. mRNA A (Gene A) bears two or more miR-146a-5p binding sites and fewer number of miRNA-2 (secondary miRNA) binding sites. Additionally, mRNA B (Gene B) harbours two or more miRNA-2 binding sites but no miR-146a-5p binding sites. **B** Possible CB regulator-target relationships considering mmu-miR-146a-5p as the regulator. Set of predicted coordinate biogenesis relationships (miRNA-1: mRNA-A→miRNA-2: mRNA-B) considering mmu-miR-146a-5p as the regulatory miRNA is exemplified here. The fold change status of differentially expressed mRNA in murine macrophages exposed to LPS (10ng/ml) for 24 hours (GSE19490) has been included in analysis to identify down-regulated ‘Gene A’ and ‘Gene-B’. **C-D** miR-146a-5p influences expression of miR-16 and miR-21-3p and their secondary effector mRNAs after LPS induction in anti-miR-146a transfected murine macrophage cells. qRT-PCR data confirms miR-16 and miR-21-3p level down regulation in anti-miR-146a transfected LPS treated RAW264.7 cells after 24 h of treatment (C). Expressions of their target mRNAs have also found to be derepressed in the same cells (D). 18s rRNA or GAPDH and U6 snRNA level were used as endogenous targets for Real time qRT-PCR based mRNA and miRNA quantification respectively. Paired two-tailed Student’s t-tests were used for all comparisons. p < 0.05 (*); p < 0.01 (**); p < 0.001(***). In (C-D) values are means from at least three biological replicates ± SD.

**Table 1:** CB regulator-target relationships likely to occur in miR-146a-5p coordinate biogenesis regulatory network in murine macrophages responding to LPS exposure.

| Gene with tandem miRNA pairs (Gene A) | CB-pair Regulator (miRNA-1) | Target gene of secondary effector miRNA (Gene B) | Secondary Effector miRNA (miRNA-2) | logFC<sup>a</sup> (miRNA-2) | P-Value<sup>b</sup> (miRNA-2) | Counts of miRNA CB-pair (prevalence) |
| --- | --- | --- | --- | --- | --- | --- |
| **bach2** | mmu-miR-146a-5p | mcm5 | mmu-miR-16-5p | 0.611762 | 0.000437 | 39 |
| **mmd** | mmu-miR-146a-5p | mcm5 | mmu-miR-16-5p | 0.611762 | 0.000437 | 39 |
| **rbl1** | mmu-miR-146a-5p | mcm5 | mmu-miR-16-5p | 0.611762 | 0.000437 | 39 |
| **irf4** | mmu-miR-146a-5p | mcm5 | mmu-miR-16-5p | 0.611762 | 0.000437 | 39 |
| **bach2** | mmu-miR-146a-5p | ncapg2 | mmu-miR-16-5p | 0.611762 | 0.000437 | 39 |
| **mmd** | mmu-miR-146a-5p | ncapg2 | mmu-miR-16-5p | 0.611762 | 0.000437 | 39 |
| **rbl1** | mmu-miR-146a-5p | ncapg2 | mmu-miR-16-5p | 0.611762 | 0.000437 | 39 |
| **irf4** | mmu-miR-146a-5p | ncapg2 | mmu-miR-16-5p | 0.611762 | 0.000437 | 39 |
| **irf4** | mmu-miR-146a-5p | mcm4 | mmu-miR-23b-3p | 0.295528 | 0.02836 | 21 |
| **bach2** | mmu-miR-146a-5p | mcm4 | mmu-miR-23b-3p | 0.295528 | 0.02836 | 21 |
| **rbl1** | mmu-miR-146a-5p | mcm4 | mmu-miR-23b-3p | 0.295528 | 0.02836 | 21 |

<sup>a</sup>logFC: Fold change in secondary effector miRNA in miR-146a over-expressing RAW264.7 cells as opposed to control.
<sup>b</sup>p-value: significance of fold change expression in secondary effector miRNA in miR-146a over-expressing RAW264.7 cells as opposed to control.

Moreover, we have studied whether CB relationships between miRNAs can have species specific differences. In order to investigate this possibility, we have predicted the probable “CB relationships” that miR-146a-5p can have in human macrophages. Herein, initially by sampling 33,214 probable CB regulator-target relationships that can occur in cells across 21 genes (Gene A) and 616 miRNA-2, a regulatory network was determined considering hsa-miR-146a-5p (miRNA-1) as the primary CB regulator. Since, expression analysis may give us an idea regarding the context dependent activation profiles of different precursor mRNA, differential expression analysis has been performed to determine which mRNA are likely to get down-regulated given that the CB regulator (miR-146a-5p) is up-regulated upon LPS exposure and it influences the biogenesis of certain secondary effector miRNA/mRNA as well. Thus, a set of 62 CB regulator-target relationships were predicted in which both ‘Gene A and B’ were down-regulated wherein there does not exist a direct regulatory relationship between ‘Gene A and B’ (Figure S2A, Table S2). Subsequently, it would have been interesting to study whether a change in the concentration of mRNA-A is reflected in the concentration of the secondary effector mRNA (mRNA-B) as we could analyse mRNA expression profiles in LPS stimulated human monocytes. Herein, we observed that miR-16 can act as a secondary effector miRNA in miR-146a-5p regulatory network responding to LPS exposure in human macrophages as well (Figure S2A, Table S2). Further, some miRNA (miR-16, miR-27b, miR-26a, miR-30e, miR-93, and miR-98) are likely to be regulated by miR-146a-5p in a similar manner in human monocytes and murine monocytes following LPS stimulation. However, there exists some species-specific differences based on the organisation of the miRNA binding sites in tandem in genes. This species-specific variation in the organization of miRNA binding sites is a possible explanation behind the species-specific variances in the networks (Figure 5B, Figure S2A). Interestingly, miR-146a-5p CB-regulatory network in macrophages may exhibit condition specific differences in regulatory network components. For instance, in macrophages responding to LPS stimuli or *Mycobacterium tuberculosis* infection a large set of different miRNAs are likely to be involved while a fraction of miRNA (miR-16, miR-20a, miR-27a, miR-27b, miR-26a, miR-22, miR424, miR-30e, miR-93, miR-96) were similar among these networks (Figure S2A, B; Table S3). Further, most of these predicted secondary effector miRNAs particularly miR-16, miR-23b, miR-26a, miR-27b, miR-30e, miR-93, miR-98 are known to modulate immune responses or inflammatory responses (Nejad et al., 2018; Prabahar and Natarajan, 2017). Therefore, by studying the CB-regulatory network of miR-146a-5p under different contexts, we determined that these miRNAs along with their secondary effector miRNAs (miR-16, miR-21, miR29b or miR-93) can potentially concordantly regulate and fine-tune macrophage inflammatory responses.

### Activation of macrophages is not prerequisite for miR-146a-5p mediated coordinated biogenesis of miRNAs

Further we wanted to explore the exclusive contribution of miR-146a-5p on biogenesis of secondary miRNAs irrespective of endotoxin-induced activation of macrophages. miR-146a-5p may have a prominent role, but thousands of different LPS-responsive molecules may also impart a crucial additive effect on miR-146a-5p mediated TDCB. To address the notion and to understand the exclusive role of miR-146a-5pon TDCB, we have adopted a miR-146a-5poverexpression system in naïve macrophage without LPS treatment. We have adopted a doxycycline-responsive (TET-ON) miR-146a-5p expressing system to validate our observation independent of macrophage activation by LPS (Figure 6A). Inducible expression of miR-146a resulted in significant upregulation of miR-125b, miR-142-3p after 24h of doxycycline induction (Figure 6B), but like our previous observations, we could not see change in the level of non-CB pair miRNA let-7a upon miR-146a-5p induction(Figure 6B). Corroborating with this, repression of endogenous targets of potential secondary miRNAs, miR-125b and miR-142-3p, could also be observed (Figure 6C). Consequently, dual luciferase reporter assay strengthened our observation further which showed increased activity of miR-125b but not of let-7a (Figure 6D) in miR-146a-5p overexpressing cells. Consistent with TDCB, cellular level of pre-miR-125b was found to be decreased post-induction correlating with increased mature miRNA processing (Figure 6E). Although miR-146a-5p has shown coordinated activity to induce specific groups of miRNAs, we wanted to check levels of different miRNPs associated proteins post-miR-146a induction to rule out the possibilities of their contribution in increased miRNP production observed for secondary miRNAs in miR-146a-5p expressing cells. However, we could not observe any significant change in expression of those factors in miR-146a-5pexpressing cells, nullifying the possibilities of miR-146a-5p mediated regulation of miRNP specific factor in modulation of the overall miRNA level inside the cells (Figure6F). FormiR-21, the other candidate miRNA that also showed the similar trend, we could see upregulation of miR-21 level and repression of its target PDCD4 after miR-146a-5p induction in macrophage cells (Figure S1G). To further rule out any other mode of cross-regulation other than miR-146a-5p mediated cooperativity in biogenesis of miR-125b, we have used the doxycycline-responsive (TET-ON) miR-146a expressing system in presence of anti-miR-146a-5pin macrophage cells to conclude on the role of miR-146a-5p in TDCB of miR-125b (Figure6G). We could see a synchronized and coordinated expression of miR-125b along with miR-146a-5p in macrophages (Figure 6H-I). We could also observe cellular level of miR-125b is connected with miR-146a abundance in macrophage even after addition of doxycycline as we could not see upsurge of miR-146a-5pactivity and miR-125b level in the miR-146a-inhibited environment (Figure6H-I). To understand the specificity of the miR-125b expression by miR-146a-5p, we could not see any changes in expression level of non-CB pair miRNAlet-7a under the same condition (Figure 6J). Altogether these data strengthen the fact that miR-146a-5p orchestrates the key steps to cooperatively modulate the biogenesis and activities of groups of secondary miRNAs in mammalian macrophage cells.

**Figure 6.**
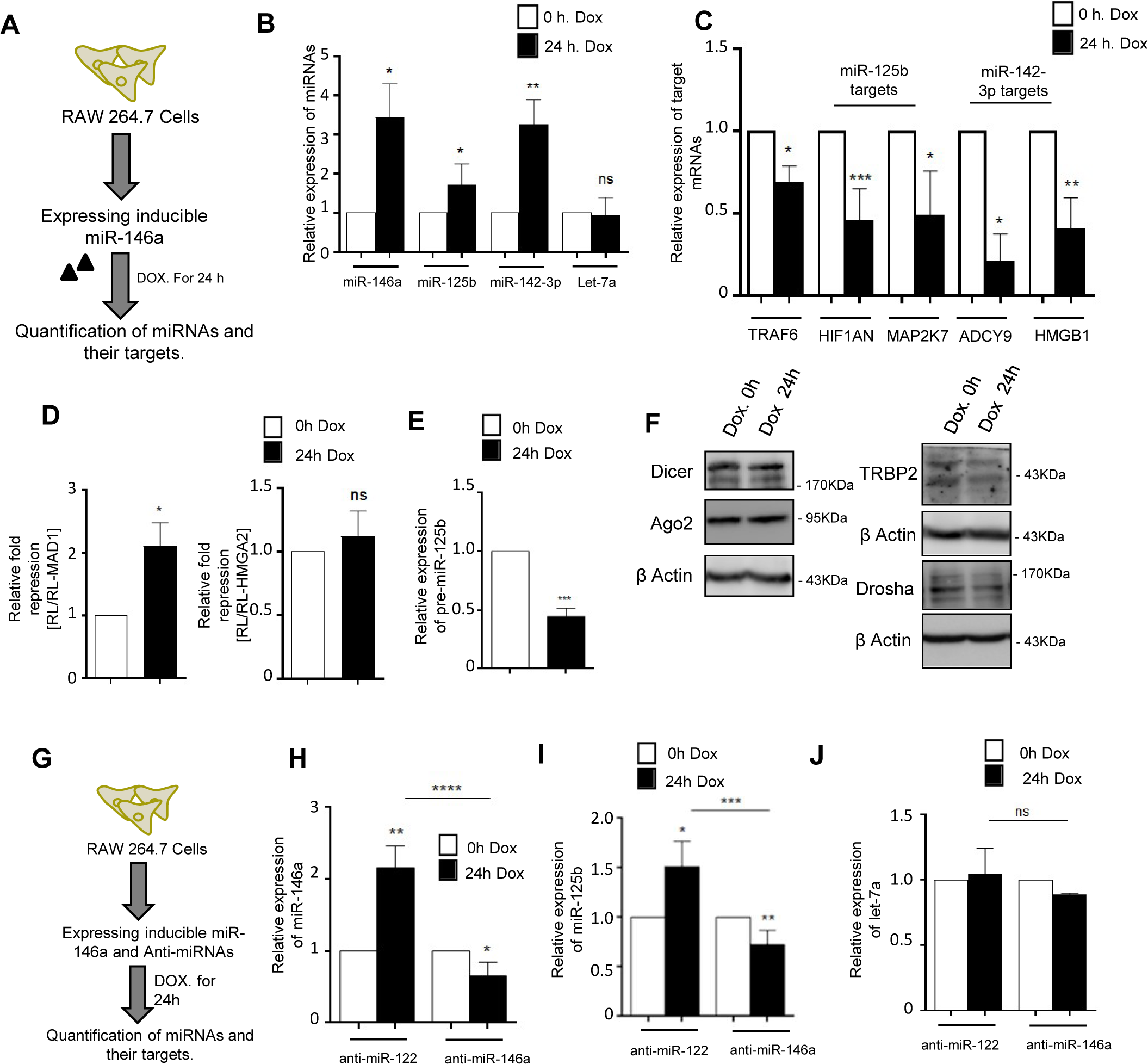
miR-146a-5p could promote coordinated biogenesis of secondary miRNAs in murine macrophage cells even without LPS-mediated activation. **A** Scheme of experiment to study miR-146a-5p mediated coordinated biogenesis of miRNAs in TET-ON RAW264.7 cells. **B** Levels of miR-125b and miR-142-3p in cells expressingmiR-146a-5p in an inducible manner in presence of doxycycline. Relative levels of miRNAs were measured after treatment of 400 ng/ml of doxycycline treatment for 24h. qRT-PCR data confirms induced expression of miR-146a-5p and coordinated upregulation of two candidate miRNAs miR-125b and miR-142-3p. Let-7a, a non-CB pair miRNA of miR-146-5p did not show any change in TET-ON RAW264.7 cells after miR-146a induction. **C** Expression of endogenous targets of miR-125b & miR-142-3p in RAW264.7 cells expressing miR-146a-5p in presence of doxycycline. The qRT-PCR data also confirmed the increased miR-146a-mediated repression of its target TRAF6 after inducible expression of miR-146a-5p. **D** Repressive activity ofmiR-125b after induction of miR-146a-5p in RAW264.7 macrophage cells. Dual luciferase assay was done with for RL-MAD1reporter mRNA having miR-125b binding sites in its 3’ UTR. RL-MAD1 but not the RL-HMGA2 that harbours seven let-7a sites on their 3’ UTR showed repression in cells expressing miR-146a-5p after 24h of doxycycline induction. RL reporters without miRNA sites were used as control. Firefly luciferase activities expressed from a construct co-transfected with RL reporters were used to luciferase activity normalization and transfection control. **E** Levels of pre-miR-125b in miR-146a-5p expressing cells after 24h of doxycycline treatment. qRT-PCR data shows significant reduction of pre-miR-125b level, suggesting increased processivity of miR-125b from its precursor form in miR-146-5p expressing cells. **F** miRNP-associated proteins do not show alteration in expression upon inducible miR-146a-5p expression in RAW264.7 macrophage cells. miRNP-associated proteins & processing enzymes Dicer1, Ago2, TRBP2 & Drosha does not show any significant change in their expression in miR-146a-5p induced cells. **G** Experimental scheme that is followed for experiments described in panel H-J where anti-miRs and doxycycline inducible miR-146a-5pconstructwere co-transfected to understand direct control of miR-146a-5p over coordinated biogenesis of secondary miRNAs in Tet-On RAW264.7 cells. **H-J** Effect of miR-146a-5p inhibition in Tet-On RAW264.7 cells expressing miR-146a-5p in an inducible manner. Reduced miR-146a-5p (H), miR-125b (I) and unchanged let-7a (J) levels in miR-146a-5p expressing cells in presence of miR-146a inhibitor but not in presence of anti-miR-122 in Tet-On RAW264.7 cells after 24h of doxycycline treatment. 18s rRNA or GAPDH and U6 snRNA level has been used as endogenous target for Real time qRT-PCR mRNA and miRNA quantification respectively. Student’s t tests were used for all comparisons. p < 0.05 (*); p < 0.01 (**); p < 0.001(***);p< 0.0001(****). In (B-E,G-J) values are means from at least three biological replicates ± SEM.

### Small RNA sequence analysis reveals coordinated biogenesis network of miR-146a-5p

The computational methodology proposed herein based on the binding site analysis provided indications regarding which miRNA (secondary effector) are likely to participate in the miR-146a-5p driven coordinate biogenesis network formation. Considering all possible combinations, it is possible that miR-146a-5p can possibly influence the expression of multiple secondary effector miRNA (187) when it acts as a miRNA biogenesis regulator considering the CB phenomenon. In order to understand whether this phenomenon can have large scale implications on the global miRNA-mRNA interaction networks within cells, we have performed a sequencing analysis to determine the likely secondary effector miRNA that have exhibited a change in their expression profile when the CB-regulator miR-146a-5p was expressed in an inducible manner. Herein, it was identified 109 miRNAs to get differentially expressed as a result of miR-146a-5poverexpression in macrophage cell lines. Among the miRNAs that got differentially expressed in the mature state, 44 miRNAs showed up-regulation while 65 miRNAs exhibited down-regulations (Table S4).

Considering this miRNA expression profiling data, we suggest that miR-146a-5p is likely to influence the biogenesis of mmu-miR-16-5p, mmu-miR-23b-3p, mmu-miR-150-5p, mmu-miR-15b-5p, mmu-miR-25-3p, mmu-miR-322-5p, mmu-miR-34a-5p, mmu-miR-500-3p. Further, we could validate the presence of secondary effector miRNA (mmu-miR-23b-3p and mmu-miR-16-5p) in the miR-146a regulatory network since these miRNAs showed significant up-regulation in their mature forms. However, some predicted secondary effector miRNA (mmu-miR-7a-5p) were found to be down-regulated as well. Additionally, considering our previous analysis of macrophage response to LPS exposure, 11 CB regulator-target relationships (Table 1) were obtained wherein miR-16-5p or miR-23b-3p (miRNA-2) biogenesis is likely to be influenced (up-regulated) by coordinated biogenesis and both primary and secondary effector mRNA (gene A, gene B) are down-regulated. It is likely that some CB-regulator target relationships could be active in particular scenarios and it would be interesting to study the validity of this phenomenon in some other contexts as well.

Subsequently, we wanted to determine the probable physiological scenarios that are likely to be driven by miR-146a-5p cooperative biogenesis network. In this respect, we identified that the validated secondary effectors like miR-16 and miR-23b are known to modulate immune responses or inflammatory responses (Nejad et al., 2018; Prabahar and Natarajan, 2017). The target genes of the probable secondary effector miRNA (mmu-miR-16-5p, mmu-miR-23b-3p, mmu-miR-150-5p, mmu-miR-15b-5p, mmu-miR-25-3p, mmu-miR-322-5p, mmu-miR-34a-5p, mmu-miR-500-3p) identified both in the sequencing analysis and the computational analysis along with predicted target genes of mmu-miR-146a-5p were considered for a pathway enrichment analysis. KEGG pathway enrichment analysis identified *Salmonella* Infection ‘mmu05132’ pathway as enriched and enriched terms herein were mainly found to occur in different signalling pathways like NF-kβ, MAPKs or apoptosis (Figure S3, Table S5).

### Coordinated biogenesis of miRNA affects cellular signalling pathways to protect cells from LPS-induced inflammation &apoptosis

LPS induction leads to activation of multiple signalling pathways like MAPKs or apoptosis. Although many of the molecular mediators are known, little is known about the miRNA-mediated crosstalk controlling these pathways. Previous reports suggest upregulation of miR-146a/b that acts as a compensatory response to curb inflammatory responses by targeting IRAK1 (of IL-1 receptor signalling), TRAF6 or decreasing TNF-α production to form a negative feedback loop (Hou et al., 2009; Nahid et al., 2009; Nejad et al., 2018) in macrophage cells. IRAK1/TRAF6 cascade signalling was known to activate p38 in a ROS dependent manner as well as phosphorylation of extracellular signal-related kinases (Matsuzawa et al., 2005; Wee et al., 2015). This intrigued us to look into differential expression of various signalling molecules of MAPKs in absence of miR-146a-5p. We could observe increased phosphorylation status of p38 and ERK1 molecules in absence of miR-146a-5p activity on LPS induced macrophages (Figure7A). In contrast, we could also observe decreased level of HSP70 protein post anti-miR-146a transfection. Activation of HSP70 through LPS was known to be involved in inhibition of NF- kβ activation thus downregulating pro-inflammatory cytokine productions (Dokladny et al., 2010; Shi et al., 2006). We have also checked the activated level of MSK1, a negative regulator of Toll-like receptor signalling, acts downstream of p38 and Erk1/2 mitogen-activated protein kinases (Ananieva et al., 2008). Decreased levels of phosphorylated MSK1 were observed in the miR-146a-5p inhibited condition (Figure7A). A significant increase of TNF-α & IL-1β levels has also been documented in absence miR-146a-5p activity-revalidating a reciprocal relationship it has with pro-inflammatory cytokines production (Figure7B). LPS induction also leads to production of iNOS expression which ultimately results in increased NO production (Lowenstein et al., 1993). Our data reveals increased generation of nitric oxide post anti-miR-146a transfection on different time-intervals after LPS induction (Figure7C). It is also evident LPS-induced TNF-α and Nitric Oxide upregulation aids in initiation of apoptosis in macrophage cells (Xaus et al., 2000). We were interested to check the level of important apoptotic marker proteins post-anti-miR-146a transfection. Apoptotic stimulation inside the cell leads to cytochrome C release from mitochondria which binds with APAF1/Procaspase 9 complex to cleave and activate Caspase 9 (Li et al., 1997). Although, post-24h & 36h of LPS induction, change in Caspase 9 was not apparent, but after 48h we could observe increased cleavage of Caspase 9 in miR-146a inactivated cells (Figure7E). Another downstream molecule of this pathway PARP, a 110 kDa protein cleaved by Caspase 3 between its Asp^214^ and Gly^215^ residue to produce the 89 kDa cleaved PARP product after the initiation of apoptosis (Boulares et al., 1999). We could observe increased cellular cleaved PARP post-anti-miR-146a transfected cells on different LPS-treatment time-point (24h, 36h, 48h) (Figure7E). Corroborating with this, TUNEL assay data shows increased apoptosis in anti-miR-146a transfected LPS-induced macrophages after 48h (Figure 7D). Our integrative observation suggests miR-146a-5p acts as crucial regulator with respect to neutralization of inflammatory responses which acts through multiple pathways; be it miRNA-mediated CB or the secondary target molecule mediated modulation of associated factors, imparts a huge role on combating pathogenic responses. This led us to propose a CB-regulated Inflammatory Network model that probably exists inside the activated macrophage cells (Fig 7F).

**Figure 7.**
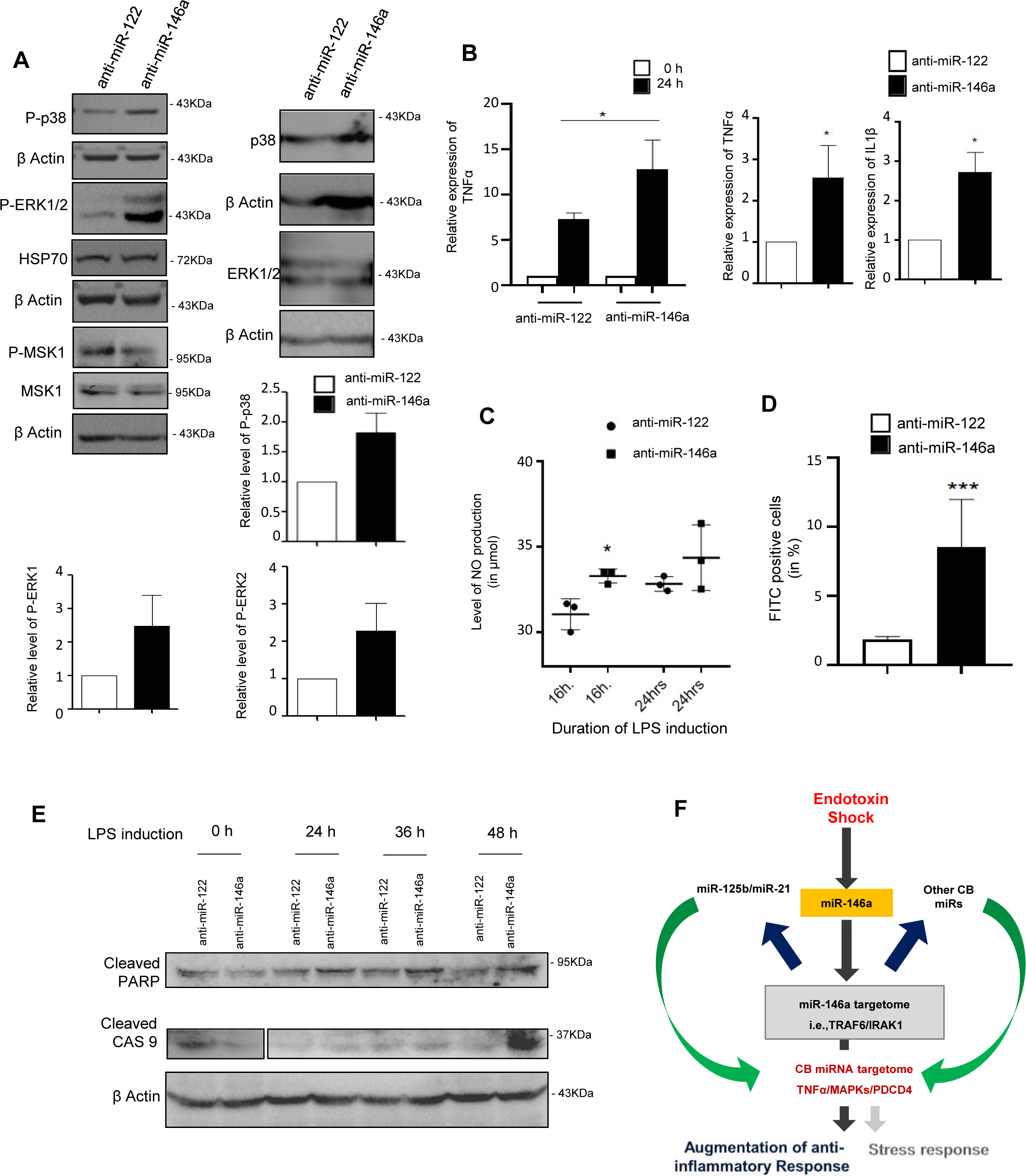
Targeting of the “Primary-miRNA” miR-146a-5p affects cellular response in LPS activated RAW264.7 cells. **A** Inhibition of miR-146a-5p activity leads to overactivation of MAPK signalling molecules. Representative western blot data confirms increased phosphorylation status of p-38 and p-ERK1/2 molecules upon LPS treatment in anti-miR-146a but not anti-miR-122 treated RAW264.7 macrophage cells. Cellular levels of HSP70 & P-MSK1 were also measured. p38, ERK1/2 and MSK1 were also detected in anti-miR-146a and control anti-miR-122 transfected cells. β-Actin serves as internal control. Comparative densitometric analysis data obtained for three such western blots were plotted to show the changed phosphorylation status of MAPK signalling molecules p-P38 and p-ERK1/2 in anti-miR146a treated cells. **B** Effect of anti-miR-146a treatment on LPS induced expression of TNF**-**α levels in RAW264.7 cells. Relative levels of proteins were measured by ELISA and plotted for both anti-miR-146a and anti-miR-122 transfected cells (left panel). Changes in levels of pro-inflammatory cytokines TNF α (middle panel) and IL1β (right panel) mRNA after 24h of LPS induction in anti-miR-146a transfected RAW 264.7 murine macrophage cells. Level in control anti-miR-122 transfected cells was considered as unit. **C** Increased generation of nitric oxide in cells with reduced miR-146a activity upon LPS-treatment. Griess assay-based quantification revealed increased NO production in anti-miR-146a transfected cells compared to controlanti-miR-122 transfected cells after 16h and 24h of LPS treatment (n=3). **D** Inhibition of miR-146a activity leads to increased induction of apoptosis of LPS treated murine macrophage cells. Percentage of apoptotic cells were calculated using TUNEL assay done for anti-miR-122 and anti-miR-146a transfected LPS induced murine macrophage cells. **E** miR-146a-5p plays a crucial role in combating LPS-induced apoptosis in RAW264.7 cells. Western blot data confirms increased cellular level of two important apoptotic marker proteins Cleaved PARP & Cleaved Caspase 9(after 48h of LPS induction) in anti-miR-146a transfected macrophage cells. **F** The probable “Coordinated Biogenesis-regulated Inflammatory Network (CBIN)” that operate inside the activated macrophage cells to balances the immune repose to protect the cells from over activation and cellular death. Blue and green arrows are representing the TDCB phenomenon and its physiological significance in murine macrophages respectively. 18s rRNA or GAPDH level has been used as endogenous target for Real time qRT-PCR mRNA quantification respectively. Concentrations and duration of LPS induction on macrophages were 10ng/ml for 24h. Student’s t-tests were used for all comparisons. p < 0.05 (*); p < 0.01 (**); p < 0.001(***). In (B-C, E) values are means from at least three biological replicates ± SEM.

### COORD-BIO web server predicts CB regulator-target relationships in humans and mice

Our previous analyses suggested that miR-146a-5p can regulate the biogenesis of specific miRNAs (particularly miR-16, miR-21, miR-142-3p, miR-125b) and the associated secondary effector mRNAs in response to LPS based stimulation of macrophage cell surface receptors. In this manner, a CB-regulator may alter the expression profile of specific miRNAs and mRNAs in turn altering or re-adjusting cellular miRNA-mRNA interaction networks as a result of the coordinate biogenesis phenomenon. Therefore, we next posed the question as to whether miRNA other than miR-146a-5p can exhibit this coordinate biogenesis phenomenon. In this respect, we have predicted the CB-regulator-target relationships for miR-155-5p and by down-regulating the miRNA and have determined whether there are corresponding changes in secondary effector miRNA that are not directly regulated by miR-155-5p. Comparative transcriptional profile of stimulated M1(LPS + IFN-γ) miR-155 knockout (KO) macrophages and unstimulated M0 KO macrophages (GSE77425) were studied. (Jablonski et al., 2016) Here, we determined that the target genes of miR-155 and target genes of secondary effector miRNA (miR-16-5p, miR-29b or miR-322) also get differentially expressed. Thus, based on our predicted CB-regulator-target relationship for miR-155, it is possible that miR-155 could potentially regulate the expression of secondary effector miRNA such as to modulate the response of macrophages upon LPS exposure (Figure S4A, Table S6). Similarly, miRNA-mRNA expression profiles in M1 and M2 polarised macrophages (Lu et al., 2016) also suggested that CB-regulator (miR-155) and secondary effector miRNA (miR-29b, miR-125a, miR-455) are simultaneously but differentially expressed there (Figure S4B, Table S6). In some CB-regulatory networks we observed that there is a possibility of a reciprocal regulation or interaction (among miR-125a and miR-155 in macrophage polarisation) as well (Figure S4C, Table S6). Therefore, different miRNA can concordantly exhibit the CB phenomenon to regulate macrophage immune response in a particular scenario. However, CB is likely to occur in different cell types as well. In this respect, we have again predicted the CB-regulator-target relationships for miR-125a-3p by considering a condition in which the miRNA had been over-expressed to determine whether there are corresponding changes in secondary effector miRNA that are not directly regulated by these miRNAs. CB-regulator-target relationships for miR-125a-3p in hematopoietic cells during cell maturation were predicted utilising a dataset (GSE33691); (Gerrits et al., 2012) wherein miR-cluster 99b/let-7e/125a had been over-expressed. It was observed that miR-125a-3p likely influences the biogenesis of miR-125b-5p, miR-338-3p, miR-706, miR-762 since ‘Gene-A’ and associated secondary effector genes (19 mRNAs as Gene B) are differentially expressed when the miR-cluster 99b/let-7e/125a is over-expressed (Table S6). These secondary effector genes are not directly regulated by the miRNA that are over-expressed but are indirectly regulated via intermediate miRNA whose expressions are likely varying as a result of the CB phenomenon. Although extensive biochemical and multifaceted approaches could confirm the existence of CB relationships for a specific scenario, it is likely that coordinate biogenesis phenomenon may exist in different physiological pathways as well to act as an additional important modulator for the fine tuning of various cellular responses.

CB-regulator-target relationships can be readily predicted based on the assumptions that we have utilised in this study. Based on the data collected during this work, we have developed a repository that can be queried to obtain a set of miRNA likely to exhibit co-ordinate biogenesis in a particular scenario since miRNA and mRNA expression data are utilised to infer the most likely relationships. It is plausible that 302 miRNAs are likely to exhibit CB phenomenon in murine cells and 1433 miRNAs are likely to exhibit CB phenomenon in human cells. Utilising these miRNA and expression profiles as query, one can obtain high confidence CB pairs wherein CB-regulator and secondary effector miRNA are up-regulated and corresponding target mRNA are down-regulated or *vice versa*. Alternately, a list of low confidence CB pairs is calculated considering that CB-regulator-target miRNA and mRNA are all expressed significantly (Figure 8A).

**Figure 8.**
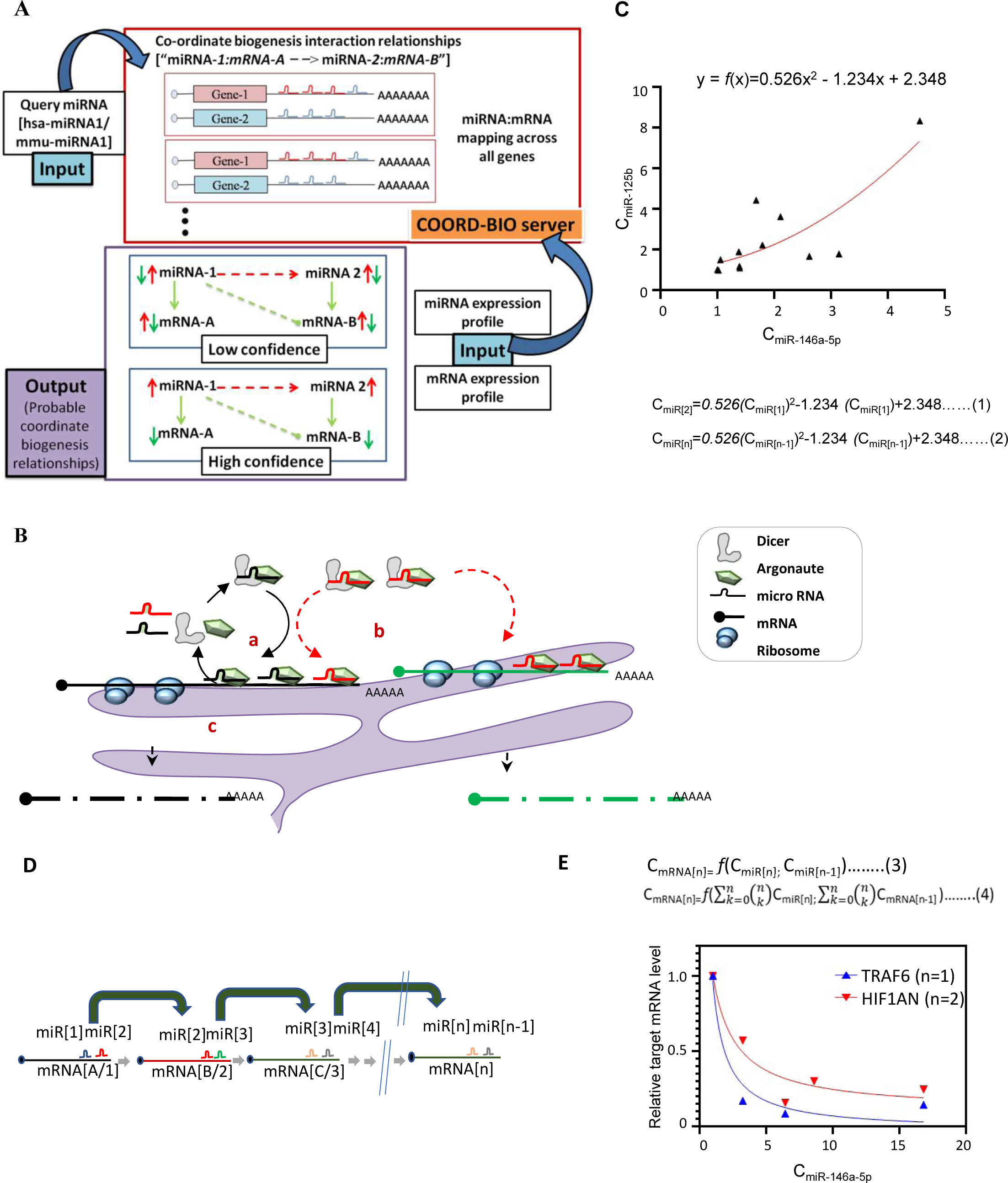
Progressive dampening mode of action of primary miRNAs on secondary targets. **A** The principal and logic gate structure for the generation of COORD-BIO web server that provides information regarding co-ordinate biogenesis relationships in *Homo sapiens* and *Mus musculus*. COORD-BIO is available at http://www.hpppi.iicb.res.in/coordB2/index.html. **B** Graphical Model of Co-operative Bio-genesis of miRNAs in Mammalian Cells. In step *a*, target mRNA (mRNA 1) dependent cognate miRNA (miRNA 1 in black) biogenesis happens and in step *b* the newly generated miRNP1 (Ago2-miRNA1) influences biogenesis and activity of other secondary miRNAs (miRNA 2 in red) that share the binding sites on the same 3’ UTR of common target mRNA in the cooperative manner. The miRNP2 (Ago2-miRNA2) targets new target mRNAs (green) that don’t have miRNA 1 target sites. In step c increased coordinated biogenesis of primary and secondary miRNAs leads to increased target cleavage or translational repression of both primary and secondary target mRNAs (mRNA A and B). **C** Change in expression of miR125b in response to changing concentration of miR146a-5p in RAW264.7 cells expressing the later miRNA in an inducible manner. The curve fitting the data point is represented by the polynomial regression equation (equation 1) considering miRNA-125b is secondary miRNA. The expression of a miRNA in a specific regulatory layer (nth layer) should be identified by proposed equation (equation 2) **D** Schematic representation of target dependent coordinated regulation of miRNA genesis showing how the concentration of miRNAs is probable getting regulated through multiple layers. **E** Probable relation of a specific mRNA concentration with the levels of its primary and secondary miRNAs (Equation 3 and 4). Plotting of data for two targets TRAF6 (primary target of miR-146a-5p) and HIF1AN (Primary target of secondary miRNA, miR-125band “coordinated biogenesis” mediated secondary target of miR-146a-5p) against primary miRNA miR-146a-5p concentration. These data suggest the dampening effect of a miRNA on their non-primary targets and the effect get reduced may be an exponential way with the distance between the subsequent layers that the miRNA and the target mRNA belong.

### Progressive dampening mode of action of primary miRNAs on secondary to tertiary miRNAs and their targets

How coordinated miRNA biogenesis affect widespread gene regulation observed in higher eukaryotes? When we plotted the concentration of miR-125b, the secondary miRNA against the changes in levels of primary miR-146a-5p in RAW246.7 cells, we found a curve connecting the data which can be presented by an equation relating changes in miR-125b level with the changes in miR-146-5p concentration (Figure 8C). It can be hypothesized for any secondary miRNAs (with lower number of binding sites on target mRNA) that the changes in its expression are primarily controlled by primary miRNA (with higher number of binding sites on the same target mRNAs) and the concentration of common target mRNA that they both targets. We have extrapolated this concept further to suggest that the concentration of the miRNA in the (n)^th^ layer in this regulatory circuit will be influenced by concentration of miRNA in the (n-1)^th^ layer and the mRNA having binding sites for (n)^th^ and (n-1)^th^ layers miRNAs. This assumption led us to formulate the equation to predict how the concentration change of individual miRNAs will affect the primary and secondary mRNAs (Figure 8D). This can be assumed that any mRNAs having binding sites for specific sets of miRNAs will not only be affected by the miRNA in the same regulatory layer (i.e in the n^th^ layer) but it would also be influenced by the concentration of miRNAs in the previous layers (n-1, n-2….etc) that are indirectly influencing the concentration of the miRNA at the same (n^th^) level through our proposed coordinated biogenesis phenomenon (Figure 8E). The distance between two layers (n and n-x where x=1,2,3…) determine the effect of a miRNA in a particular layer on the mRNA targets in another layer and the repressive effect should decrease gradually with the value of x. Therefore, the expression of a mRNA is influenced by sum of the effects of individual miRNAs present in different layers that do not require to have one canonical binding site on that mRNA. Thus, the miRNA repository of a cell is influenced by the status of multiple miRNAs and the network they form which have the cumulative effect on the mRNA transcriptome of the cell (Figure 8D, E). A disease condition that affects specific primary miRNA pool thus could be connected with changed expression of the “secondary” target genes which are without the binding sites for the primary miRNAs but are differentially expressed prominently in disease condition due to changed levels in the secondary or tertiary miRNAs.

## Discussion

Here we report Target Dependent Coordinated Biogenesis (TDCB) of miRNAs where target-mRNAs increased-biogenesis of a family of secondary miRNAs that share the binding sites in the 3’ UTR of common target mRNAs having binding sites for a primary miRNA. We have identified and unravelled how the TDCB operates in a physiological context to regulate miRNA-network and fine-tune the inflammatory responses. The previous works have been centred upon regulation of a single miRNA activity by its targets, our findings emphasized on understanding of the sequential molecular events in a physiological context that drives cascaded expression of miRNAs affecting their expression at post-transcriptional level. Earlier PDCD4 mediated co-regulation of miR-21 and miR-499 has been documented, where activity and stabilization of one miRNA species has been shown to influence others (Ajuyah et al., 2019). We have proposed the coordinated biogenesis model of miRNA where presence of multiple target sites of a primary miRNA increases the fall off rate of Dicer1 molecule from the miRISC for its reassociation with new Ago2 to form new miRISC with secondary miRNAs and favour their association of shared 3’ UTR. This, in turn, not only increases the biogenesis of primary miRNA but also other secondary miRNAs having adjacent targets sites on the coregulated mRNAs in a coordinated manner. These results in repression of secondary mRNAs targeted by these newly generated secondary miRNAs but not by the primary miRNAs (Figure 8B). These findings broaden our basic understandings from a “one miRNA to one mRNA’’ regulation to miRNA-targetome based coordinated gene repression network in the field of miRNA-mediated gene regulation where cooperativity among target sites influences the biological processes.

Importance of membranes of mammalian cells have been documented as the primary sites for miRNA-mediated processes including the target dependent microRNA biogenesis (Bose et al., 2020). Here we have documented CB of miRNA occurs on the rER membrane when the primary target mRNA helps the compartmentalization and biogenesis of both CB-pair miRNAs and in turn helps in compartmentalization of the secondary target mRNA to the polysomes attached with the rER. This helps the recruitment of secondary messages that otherwise not part of the protein translation regulating machineries attached with rER. CB, in turn, thus contributes to the gene regulation process by mRNA compartmentalization-suggesting a new paradigm in gene regulation and mRNA trafficking.

We hypothesized TDCB is a general and complex phenomenon probably occurring on every other miRNA-driven pathway but it was challenging to single out the phenomenon due to existence of a subtle crosstalk of different interlinked pathways inside the cell. We were looking for a well-conserved miRNA responsive pathway where miRNAs act prominently to regulate multiple pathways, making it a suitable candidate to showcase TDCB. Expressions of many endotoxin-responsive miRNAs are mostly controlled in a NF-kβ dependent manner, their mode of action varies as they form reciprocal regulatory relationships also. Like, miR-155 and miR-146a-5p have been proven to be the critical reciprocal mediators of this phenomenon (Huffaker et al., 2012; Mann et al., 2017). NF-kβ binds to the putative promoter elements for many endotoxin responsive miRNAs in a time-dependent manner that diversifies the response as immediate early, early and later immune responses (O’Neill et al., 2011; Zhou et al., 2010). This led us to speculate that there might be an existence of sensing mechanisms amongst miRNAs that coordinates the diversified network to balance the immune responses inside the macrophages. This intrigues us to unravel the molecular mediators of the phenomenon. Absence of a prominent anti-inflammatory miRNA, miR-146a-5p on LPS induced macrophage cells shows differential expression of LPS-responsive miRNAs that share target sites with miR-146a-5ptargetome. To broaden our observation, whole cell transcriptomics data also suggests a group of miRNAs also exhibits miR-146a mediated CB, irrespective of pre-documented LPS induced behaviour. An integrated biochemical and computational approach has been designed to predict and validate probable regulatory relationships between miRNAs wherein one miRNA is likely to influence the biogenesis of the other. As per our hypothesis, a CB regulator (miRNA-1) is up-regulated to induce biogenesis of another miRNA (miRNA-2) by coordinated action while the corresponding targets (mRNA-A) with binding sites for both miRNAs are likely to be down-regulated. Further, as a result of this phenomenon, a corresponding downstream effect on a class of secondary mRNA targets (mRNA-B) containing the binding sites for the secondary effector miRNA-2 should be observed. In order to study this mechanism, we have utilized miRNA-mRNA regulatory interaction data to predict a set of miRNA which are likely to act as CB regulator and influence the biogenesis of another miRNA target. Thus, a probable miRNA biogenesis regulatory or meta-interaction network has been predicted in mice and humans. Utilizing large scale analysis of expression data of monocyte exposed to LPS, we could identify a number of miRNAs which are likely to be regulated by miR-146a-5p. miRNAs such as miR-16-5p, miR-26a-5p, miR-27b-3p, miR-30e-5p, miR-93-5p, miR-98-5p among those which have been identified as probable secondary miRNAs in miR-146a-5p regulatory network of coordinate biogenesis in mice and humans. Moreover, the predicted CB relationships (miRNA-1:mRNA-A to miRNA-2:mRNA-B) for mmu-miR-146a-5p have been studied in detail and we could validate that miR-146a-5p likely regulates the biogenesis of miR-16-5p, miR-23b-3p, and miR-21-3p in turn regulating the expression of secondary effector mRNA as well. By studying the miRNA binding site location and number, we could predict possible CB pairs and their network of secondary effector miRNA and mRNA. However, there are subtle differences between the predicted and the experimentally analysed dataset, which could be due to the lack of granularity of the sequencing data where all binding sites may not have been characterized. Further, prevalence (counts) of different miRNA pairings (CB relationship pairs) that occurred across all Gene A, were determined. Some of these relationships along with others that had literature support were studied in detail. A substantial number were studied to establish our proposed hypothesis. We have also utilised other large scale expression analysis datasets to support the TDCB paradigm. miR-146a-5p have been identified to exhibit this co-ordinate biogenesis phenomenon to regulate macrophage immune response upon stimulation via lipopolysaccharide.

Existence of different endotoxin-responsive signalling modulators of NF-kβ, MAPKs, Akt1signalling pathways and endotoxin-responsive miRNAs interacts amongst each other to create a crosstalk to elicit the cascaded immune responses. Hence to negate the influence of endotoxin-responsive molecules on miR-146a-5p, we have adopted an LPS-free miR-146a overexpression system. Precisely inducing the miR-146a expression on macrophages and monitor the after-effect related to coordinated miRNA biogenesis and function. Biochemical analysis and RNA sequencing data corroborates with each other that strengthens and revalidated our observation found on activated macrophages also.

On a precise note, these mechanisms probably have finer levels of cooperative regulation vested upon multiple co-mediators, that this may not be exclusively controlled by groups of TDCB class of miRNAs. Rather the integrative network formed by the “miRNA” and “effector target mRNAs/proteins regulated by miRNA” imparts the highest level of cooperation to elicit this response. Ultimately this inter-connected complex network gives rise to a signalling web where CB is probably eliciting the phenotypes. For instances, regulation of TNF-α expression might be controlled at different levels by different modulators. IRAK1/TRAF6-controlled phosphorylation of p38 & ERKs activates different MAPK-activated protein kinases to further activate different transcription factors, production of pro-inflammatory cytokines, and cell surface receptors that may ultimately lead to onset of cellular apoptosis(Chi et al., 2006; Chuang et al., 2000; Roux and Blenis, 2004). Increased production of TNF-α attribute a huge impact on iNOS synthesis also (Chan and Riches, 2001; Chen and Wang, 1999). TNF-α also bears binding sites for miR-125b (Tili et al., 2007). Probably this layers of cascaded regulation manifests in the temporal expression of TNF-α. Contrastingly for another instance, miR-21, another important CB candidate has a significant repressive effect on p38-CHOP and JNK signalling to inhibit pro-inflammatory phenotype and macrophage apoptosis modulated by MAP2K3 expression (Canfran-Duque et al., 2017). PDCD4, another pro-inflammatory protein is also suppressed by miR-21 that leads to decreased NF-kβ activity and increased production of anti-inflammatory cytokines IL-10 (Sheedy et al., 2010). We could observe significant derepression of these molecules (i.e.MAP2K3 & PDCD4) on miR-146a-5p inhibited conditions also, wherein both of these mRNAs does not bear sites for miR-146a. This “miR-146a→TRAF6→miR-21→PDCD4/MAP2K3/PTEN→p38/NF-kβ/JNK” mediated probable regulatory network axis might be one among many that exist in activated macrophage. These findings led us to predict Coordinated Biogenesis-regulated Inflammatory Network (CBIN) (CBIN) that probably exists in cells post-endotoxin treatment. There is other multi-linear co-regulation that has been also reported on inflammation context that even further supports our TDCB-based interactive analysis for fine-tuning each and every orchestrated response (Kim et al., 2012; Rajaram et al., 2011). Altogether, we could see groups of familiar miRNAs acts in coordination but in an target-dependent manner to protect the macrophage cells through pro-inflammatory cytokine silencing, inhibition of apoptotic pathways resulting in tissue healing.

Along with we have also determined the likely conditions under which this CB phenomenon can occur and developed a computational module to predict regulatory miRNA and their network under these particular contexts. Analysis of few CB regulatory networks suggests that multiple miRNA may potentially exhibit this phenomenon in a condition specific, cell type specific and species specific manner. There might be also other different factors which influence our observation. Stabilization and turnover of miRNA, strength of miRNA-target site pairing, distance between cooperative sites or influences of other RNA binding proteins may be proven crucial for this type of cooperation. Not only Dicer processivity or complementary seed sequence pairing, spatial binding among molecules might also impart a crucial role for this type of interaction. Further experiments on multiple various aspects may give a better understanding of the structural basis of this phenomenon. Overall, we have unravelled a unique phenomenon where TDCB of miRNAs contributes to build a cascaded molecular framework to balance the immune responses in activated macrophages.

## Materials & Methods

### Cell Culture and Reagents

TET-ON stable HEK293 cells with the Tetracycline-inducible expression constructs were grown in DMEM supplemented with 10% TET-approved FCS (Clontech) and induction of expression of respective gene product was carried out for indicated time durations using 400 ng/ml Doxycycline (SIGMA). HEK & HeLa cells were maintained in DMEM medium containing 2 mM L-glutamine and 10% heat-inactivated FCS as described earlier (Bose and Bhattacharyya, 2016). RAW 264.7 cells were cultured in RPMI 1640 medium (Gibco) supplemented with 2 mM L-glutamine, 0.5% β-Mercaptoethanol and 10% heat-inactivated fetal calf serum (or 10% TET-approved FCS for experiments related to tetracycline-inducible expression constructs system). Primary murine peritoneal exudates macrophages (PEC) were boosted by 4% starch injected intra-peritoneally. After 72 hrs. of starch mediated stimulation, mice were sacrificed and peritoneal macrophages were collected after washing and clearing of the peritoneal cavity with cold 1X PBS. Cells were spun down and grown in RPMI 1640 supplemented with 2 mM L-glutamine and 10% heat-inactivated fetal calf serum (FCS).

### Expression Plasmids, Transfection & Treatment on Cells

Plasmids bearing humanized renilla luciferase coding sequences (RL-con), three let-7a binding sites downstream of the renilla luciferase (RL) coding region (RL-3xbulge-let-7a), and firefly luciferase (FL) under a simian virus 40 (SV40) promoter (pGL3FF) were kind gifts from Witold Filipowicz. Experiments related to RL-WT(RL-3xbulge-let-7a_1xbulge-miR-122) or Mut.(RL-3xbulge-let-7a-mut_1xbulge-miR-122) constructs, 1xbulge-miR-122 sequence cloned downstream of 3xbulge-let-7a(WT) or 3xbulge-let-7a_mut(with mutation on seed sequence) sequence in NotI site of pRL-3xbulge-let-7a. Regarding further study related to secondary mRNA expression, RL-sequence has been replaced with GFP sequence to make the GFP-WT (GFP-3xbulge-let-7a_1xbulge-miR-122) or GFP-Mut (GFP-3xbulge-let-7a-mut_1xbulge-miR-122) constructs. Pre-miR-122 or pre-miR-146a sequence cloned within pTRE-Tight-BI Vector (Clontech) in NheI and NotI sites or Bam HI and Hind III to make induced miR-122 (imiR-122) and induced miR-146a (imiR-146a) construct respectively. pTet-On® Advanced Vector (Clontech) expressing Tet-responsive reverse transactivator were used to activate Tet-On system inside the cells. Plasmid expressing GFP protein amplified and cloned in pCI-neo vector to make GFP-Con. Precursor miRNA-125b sequence cloned into pU61 Hygro (Genescript, USA) vector and MAD1-3′UTR was cloned into the linearized pMIR-REPORT vector (Applied Biosystems, Foster City, CA, USA) downstream of the Luc gene to make pmiR-125b and pRL-MAD1-3’UTR respectively, were kind gifts from Dr.Susanta Roy Chowdhury (Bhattacharjya et al., 2013)). FLAG and HA tagged human AGO2 expression plasmid FHA-AGO2 were kind gifts from Gunter Meister (Meister et al., 2004). The Human IRAK1-3′UTR was cloned into downstream of pRL-con vector using the XBaI and NotI restriction sites to make pRL-IRAK1-3’UTR construct. Downstream of the pRL-con plasmid 3’ UTR of the HMGA2 gene with intact let-7a binding sites were cloned (RL-HMGA2 3’UTR) that are a kind gift from Anindya Dutta (Barman and Bhattacharyya, 2015).

1 μg target mRNA encoding plasmids (WT or Mut constructs) were transfected in six-well format with 500 ng inducible miR-122 encoding plasmid on Tet-ON HEK stable cell lines. For GFP-3 × bulge-122 reporter expression study on HEK293 cells, exactly the same transfection were performed with addition of this plasmid. Lipofectamine 2000 (Life Technologies) has been used as transfection reagent for all of this study as per the manufactures protocol. 30 nM of Ambion® Anti-miR™ miRNA Inhibitor, either anti-miR-146a-5p (AM10722), anti-miR-155-5p (AM13058) or anti-miR-122-5p as control (AM11012) was transfected on RAW 264.7 cells for anti-miR experiments. The macrophage cells were induced with 10 ng/ml Escherichia coli O111:B4 LPS (Calbiochem, La Jolla, CA) for activation unless specified otherwise. For immunoprecipitation study, 1.5 μg of FLAG-HA-AGO2 expressing plasmid has been transfected along with anti-miRs on 6 well formats. Transfection method has been followed as per the manufacturer’s protocol. siDicer1 specific for mouse cells were transfected at 30 nM concentration on RAW 264.7 cells. Dharmacon SMARTpool ON-TARGETplus siRNAs against Dicer1 were bought from Thermo Scientific. siCon (ON-TARGETplus non-targeting pool) has been used as an control. Transfection of siRNAs has been carried out using RNAiMax (Life Technologies) as per the manufacturer’s protocol. Experiments related to inducible expression systems on RAW 264.7 cell, 600 ng of Tet-On plasmid and 400 ng of inducible miR-146a plasmid has been transfected on 12 well cell culture formats. Fugene HD (Promega) transfection reagent has been used for this as per manufactures protocol.

### Luciferase assay

Luciferase reporter assays were carried out using Dual-Luciferase Assay Kit (Promega), following the manufacturer’s instructions, on VICTOR X3 Plate Reader (PerkinElmer). Renilla luciferase (RL) luminescence has been normalized with Firefly luciferase (FF) activities to calculate fold repression. In the experiment using RL target reporters, HEK293 cells were transfected in a 24-well format with 250 ng imiR-122, 200 ng firefly luciferase, along with 20 ng of Renilla luciferase reporter RL-3xbulge-let7_1xbulge-miR-122 (WT or Mut). After 24 h cells 400ng/ml of Doxycycline has been treated and at 48 h Luciferase activity measured. Firefly normalized RL values were plotted. For Luciferase reporter study on RAW cells, 200 ng Firefly Luciferase (FL), along with 40 ng of Renilla Luciferase (RL) reporter has been transfected along with necessary reporter plasmid or anti-miRs.

### Animal Ethics

The animal facility of CSIR-Indian Institute of Chemical Biology has provided all the required Mice Adult Balb/C mice (of any sex) and we have followed the National Regulatory Guidelines issued by the Committee for the Purpose of Supervision of Experiments on Animals, Ministry of Environment and Forest, Govt. of India for all the experimentations we have performed.

### Isolation of RNAs, Real-Time PCR quantification of mRNA and miRNAs

TRIzol reagent (Life Technologies) have been used for isolation of RNA from total cellular level or from organelle fractions. For mRNA quantification, random nonamers (Eurogentec Reverse Transcriptase Core Kit) was used to prepare cDNA by using the 200 ng of the total RNA for 10 μl of reaction and the produced cDNA was used for comparative quantitation of mRNA expression. Real-time (reverse transcriptase) PCR from cDNA was done with the Mesa Green qPCR Mastermix Plus for SYBR Assay-Low ROX (Eurogentec). For real time based quantification to study comparative differential expression here are list of primers that has been used: CAT1 (Forward: 5’ GCCGCCGGCTTGGATTCTGA 3’, Reverse: 5’ CCCCGAGGGCCACCAGATCA 3’), ALDO A (Forward: 5’ TGGACCTAGCTTGGCGCGGA 3’, Reverse: 5’ CCTGGGCCAGCAGGCAGTTC 3’), GYS-1 (Forward: 5’ GGTGGCTAACAAGGTGGGTGGC 3’, Reverse: 5’ CGATCAGCCAGCGCCCGAAA 3’), N-RAS (Forward: 5’ TCATGGCGGTTCCGGGGTCT 3’, Reverse: 5’ TCAACACCCTGTCTGGTCTTGGC 3’), KRAS(Forward: 5’ GCAAAGACAAGACAGAGAGTGGAGG 3’, Reverse: 5’ AACTGCATGCACCAAAAACCCCA3’), TRAF6 (Forward: 5’ CCTCAAGATGTCTCAGTTCCATC 3’, Reverse: 5’ GTTCTGCAAAGCCTG CATC 3’), IRAK1 (Forward: 5’ CTGTGGCCCTGGATCAAC 3’, Reverse: 5’ GAAAAGCTGGGGAGAGGAAG 3’), HIF1AN (Forward: 5’ AGTGCCAGCACCCATAAGTTC 3’, Reverse: 5’ AACCCAAGAAG TCCATGACAATC 3’), MAP2K7 (Forward: 5’GATCCCACCAAGCCTGACTATG3’, Reverse: 5’ACTGGAAGTCCCCTGAGAAGCC3’), HMGB1 (Forward: 5’ GGCCTTCTTCTTGTTCTGTT 3’, Reverse: 5’GCAACATCACCAATGGATAA 3’), ADCY9 (Forward: 5’ GCAAAATGGCTGTCAA GACGAGC3’, Reverse: 5’ CTGGCTGTTAGTGAGCTTCTCC 3’), PDCD4 (Forward: 5’ GTAGATTGTGTACAGGCTCGAG3’, Reverse: 5’ CCCACACACTGTCTTTCCGC 3’), MAP2K3 (Forward: 5’ GAGGCTGATGACTTGGTGAC3’, Reverse: 5’ GCACCATAGAAGGTGACAGTG 3’), Pre-miR-155 (Forward: 5’ CTGTTAATGCTAATTGTGATAGG3’, Reverse: 5’ CTGTTAA TGCTAACAGGTAGG 3’), Pre-miR-146a (Forward: 5’ AGCTCTGAGAACTGAATTCC3’, Reverse: 5’ GCTGAAGAACTGAATTTCACAG 3’), Pre-miR-21 (Forward: 5’ TGTCGGATAG CTTATCAGACTG3’, Reverse: 5’ TGTCAGACAGCCCATCG 3’), Pre-miR-125b (Forward: 5’ AGTCCCTGAGACCCTAACTTG3’, Reverse: 5’ AGCTCCCAAGAGCCTAAC 3’), Mis18bp1 (Forward: 5’ TCTTCCAAAGCACAAACCTGG3’, Reverse: 5’ TTCCCACTTTGGCAGTTATC 3’), MCM 5 (Forward: 5’ ATCCAGGTCATGCTCAAGTC3’, Reverse: 5’ TTCCTGGGAAGGGCATAGC 3’), NCAPG2 (Forward: 5’ ACTAGATGAATTATCAAGGAAAC3’, Reverse: 5’ CATTTATT ATAGATACAGAAGCAAG 3’), RRM2 (Forward: 5’ TTTCTATGGCTTCCAAATTGC3’, Reverse: 5’ GATGCAAAAGAACCGGAAAAG 3’), ASPM (Forward: 5’ ATATTAACCCCTGATGACTTC3’, Reverse: 5’ AACCATTTTTTCAGAAGTAAAC 3’), NCAPD3 (Forward: 5’ AAGGCTT TTCATATCTGGTCC3’, Reverse: 5’ TTGGGAAGATGCTTTGCAATATG 3’), FEN1 (Forward: 5’ AACACAATGATGAGTGCAAAC3’, Reverse: 5’ ATGCACAGATCCACAAACTG 3’), TNF α (Forward: 5’ GTCTCAGCCTCTTCTCATTCC3’, Reverse: 5’ TCCACTTGGTGGTTTGCTA 3’), IL 1β (Forward: 5’ GACCTTCCAGGATGAGGACA3’, Reverse: 5’ CCTTGTACAA AGCTCATGGAG 3’), GFP (Forward: 5’ TGACCACCCTGACCTACGG3’, Reverse: 5’ GAAGTC GATGCCCTTCAGC 3’). 18s rRNA (Forward: 5’ TGACTCTAGATAACCTCGGG 3’, Reverse:5’ GACTCATTCCAATTACAGGG 3’) or GAPDH (Forward: 5’ CAGGGGGGAGCCAAAAGGG 3’, Reverse:5’ CTTGGCCAGGGGTGCTAAGC 3’) has been used as endogenous control. The comparative Ct method, have been used for relative quantitation that typically use normalization by the housekeeping mRNA expression viz. 18S rRNA or GAPDH levels.

Cellular or organellar miRNA levels were quantified using TaqMan-based miRNA specific assay kit starting with 50 ng of total RNA. The reverse transcription mix was used for primary PCR amplification of miRNAs and finally with TaqMan Universal PCR Master Mix No AmpErase (Applied Biosystems). The samples were probed in triplicates for each biological replicates. The comparative Ct method has been used where the candidate miRNA has been normalized with endogenous control U6 snRNA. The candidate miRNA of whose specific primers were used are for let-7a (assay ID 000377), miR-122 (assay ID 000445), miR-146a (assay ID 000468), miR-155 (assay ID 002571), miR-21 (assay ID 000397), miR-125b (assay ID 000449), miR-142-3p (assay ID 000464), miR-16 (assay ID 000391), miR-21-3p (assay ID 002493) and U6 snRNA (assay ID 001973). All reactions were performed in 7500 Applied Biosystem Real Time System or BIO-RAD CFX96 Real-Time system. Cycles were set as per the manufacturer’s protocol.

### Immunoprecipitation

Procedure for immunoprecipitation of Ago2 was followed as per published protocols ((Bhattacharyya et al., 2006; Kundu et al., 2012). FLAG-M2 agarose beads (Sigma) were washed thrice with 1× IP buffer (20 mM Tris–HCl, pH7.5; 150 mM KCl, 1 mM MgCl_2_) for exogenous FLAG-tagged Ago2. Cells were lysed in a lysis buffer [20 mM Tris-HCl pH 7.4, 200 mM KCl, 5 mM MgCl_2_, 1 mM dithiothreitol (DTT), 1× EDTA-free protease inhibitor (Roche), 5 mM Vanadyl ribonucleoside comples (Sigma), 0.5% Triton X-100, 0.5% sodium deoxycholate] at 4 °C for 30 min, followed by 10-s sonication for thrice with 5-min incubation on ice in between. Lysate was centrifuged at 16,000 × g for 5 mins. at 4°C after lysis. Immunoprecipitation was carried out for 16 h at 4 °C. After washing the beads with 1X IP buffer, beads were divided in two halves for protein and RNA expression analysis. One half subjected to RNA isolation with TRIzol LS and another for western blotting with 1× SDS sample buffer.

### Isolation of cellular membranes, microsomes and polysomes

50 µg/ml digitonin (Calbiochem) has been treated to the cells for 10 min at 4°C followed by centrifugation at 2,500*xg* as described (Barman and Bhattacharyya, 2015). Isolation of Polysome and Microsome were performed as elaborated in detail earlier (Chatterjee et al., 2020). Following isolation of respective cellular fractions, equivalent amount of RNA was used to check organeller enrichment of miRNAs and their respective target mRNAs over total cellular fractions.

### Western Blotting

Western blotting of proteins has been followed as per published protocols (16081698). The cell lysates or immunoprecipitated proteins were electrophorated through SDS–PAGE. After completion of electrophoresis, proteins were transferred to polyvinylidene fluoride or polyvinylidene difluoride (PVDF) membrane followed by blocking at 4°C for minimum 1 h. The blots were probed for overnight at 4°C with the following sets of primary antibodies: mouse anti-Ago2 (Abnova), 1:1000; rabbit anti-DICER1 (Bethyl) 1:8000; rat anti-HA (Roche), 1:1000; HRP-conjugated anti-β-Actin (SIGMA), 1:10000; rabbit anti-TRBP2 (Cell Signalling), 1:1000; rabbit anti-Drosha (Bethyl), 1:8000; rabbit P-p38 (Cell Signalling), 1:1000; rabbit anti-P-ERK1/2 (Cell Signalling), 1:1000; rabbit anti-P-MSK1 (Cell Signalling), 1:1000; rabbit anti-MSK1 (Cell Signalling), 1:1000; rabbit anti-HSP70 (Cell Signalling), 1:1000; rabbit anti-Cleaved PARP (Cell Signalling), 1:1000; rabbit anti-Cleaved Caspase 9 (Cell Signalling), 1:1000; rabbit anti-P-Akt (Ser-473) (Cell Signalling), 1:1000; Visualization of all Western blots was performed using an UVP BioImager 600 system equipped with VisionWorks Life Science software (UVP) V6.80. ImageJ software has been used for densitometric analysis of blots for relative quantification of bands.

### Prediction of coordinate biogenesis interaction relationship data

Experimentally determined miRNA binding site information (genomic location, number of sites) was collected from TarBase (Vlachos et al., 2015) and miRTarBase (Chou et al., 2016) database. Interactions (2,33,601) between human miRNA (2588) and their target genes (8808) from miRTarBase (both ‘Strong’ and ‘Weak’ interactions, which had genomic binding positions listed were considered) were retrieved. Similarly, 24,650 interactions were retrieved between mouse miRNA (847) and their target genes (3940) as well. Additional interactions (2,82,181 and 1,34,997, respectively) were retrieved between human miRNA (1026) and target genes (13,811), and mouse miRNA (434) and genes (10,581) from the TarBase (‘Direct’ interactions which had genomic binding positions listed were considered) (Vlachos et al., 2015). TarBase and miRTarBase data were combined and pooled together to extract all possible experimentally verified miRNA-mRNA interactions in human and mouse, respectively.

Under certain conditions miRNA can influence the biogenesis of other miRNA, a phenomenon referred to as coordinate miRNA biogenesis. In order to computationally predict miRNA likely to exhibit coordinate biogenesis, CB regulator and its associated targets were identified according to the following conditions:

Condition 1: miRNA-1 (eg: miR-146a) should have 2 or more binding sites in tandem with miRNA-2 (with fewer number of binding sites than miRNA-1) on the same 3’ UTR of the common target gene/mRNA (within any gene say ‘Gene A’).

Condition 2: Target gene (‘Gene B’) of secondary effector miRNA (miRNA-2) should contain 2 or more binding sites of miRNA-2 and does not carry binding sites for miRNA-1 on its 3’ UTR.

These two conditions generated multiple triads each of which includes ‘miRNA-1’, ‘miRNA-2’ (secondary effector) and ‘Gene B’ (secondary effector gene) (Figure 4A). Further, the expression level of ‘secondary effector gene’ may indicate change in miRNA-2 biogenesis (secondary effector) mediated by miRNA-1. “miR-146a-5p” is the miRNA-1 in our context as we wanted to explore probable cooperative effect of this miRNA.

This relationship [“miRNA-1:mRNA-A --˃ miRNA-2:mRNA-B”] derived between miRNA-1 and miRNA-2 considering the binding site information data has been utilized to determine the coordinate biogenesis regulatory network for miR-146a-5p in *Mus musculus* (Table S7) and *Homo sapiens*, respectively. Additionally, a prevalence count corresponding to different miRNA coordinate biogenesis pairs that occur across all ‘Gene A’ has been determined; this value can be utilized to assign precedence to more likely coordinate biogenesis pairs. Further, in order to determine a set of regulator-target relationships that we can validate or study in an experimental set up, we have utilized large scale expression analysis datasets to prune our regulator-target relationships list.

### Contextualization of coordinate biogenesis pairs for experimental validation

Expression analysis data from murine macrophages and isolated human monocytes that were transformed into macrophages exposed to LPS (10ng/ml) for 24 hours (similar condition as experimental datasets considered in Figure2) were considered [GSE19490, (Schroder et al., 2012); GSE85333, (Regan et al., 2018)]. Differentially expressed mRNAs in each case were determined considering a fold change threshold of 1.5 and p-value threshold of 0.05. Further, these differential expression data were mapped in Homo sapiens and Mus musculus coordinate biogenesis regulator-target relationship data for miR-146a-5p according to the following conditions.

Regulator-target relationships in which secondary effector gene (Gene B) and primary target for miR-146a (Gene A) were both found to be down-regulated, considering “miR-146a mediated coordinated repression” were selected. The respective fold change and p-value have been included in Table S1. Moreover, coordinate biogenesis miRNA pairs occurring in genes where ‘Gene A’ has a direct regulatory relationship with ‘Gene B’, for instance, ‘Gene A’ could regulate ‘Gene B’ as a transcription factor have been filtered out from the validation set. The filtering was performed to ensure the effect on the secondary effector gene is most likely due to the increase in biogenesis of miRNA-2 as a result of the coordinate biogenesis relationship that they are likely to share.

### Validation of CB regulator-target pairs utilizing large scale expression profiling

Small RNA sequencing analysis to determine differentially expressed miRNA in the background of miR-146a over-expression in RAW264.7 cells had been performed to identify miRNAs whose expression are influenced when the CB regulator (miR-146a) expression is up-regulated. miRNA that had fold change (F.C.) > 0 and p-value <= 0.05 have been considered as up-regulated and miRNA that had fold change (F.C.) < 0 and p-value <= 0.05 have been considered as down-regulated. Since CB regulator expression is up-regulated I should in turn up-regulate the expression of another miRNA (miRNA-2) by co-ordinate biogenesis. Thus, utilizing miRNA expression profiles such CB-regulator target relationships for miR-146a-5p have been identified. It has further been studied whether these CB regulator-target relationships are widely observed in macrophages responding to LPS exposure wherein miR-146a is likely to be over-expressed. Additionally, considering target genes of miR-146a and secondary effector miRNA in miR-146a coordinate biogenesis network, a pathway over-representation analysis (Yu et al., 2012) was performed to determine the physiological relevance of this network. Further, similar CB regulator-target relationships have been identified for other miRNA likely to be over-expressed in macrophages as a result of LPS exposure and these have been further validated with the help of joint miRNA-mRNA expression profiling data.

### Determining co-ordinate biogenesis regulatory network for miR-155-5p, miR-125a-3p and miR-146a-5p

The co-ordinate biogenesis relationship [“miRNA-1:mRNA-A --˃ miRNA-2:mRNA-B”] were determined utilizing the conditions (condition 1 and 2) mentioned above by considering mmu-miR-155-5p as miRNA-1 to determine the probable coordinate biogenesis regulatory network of miR-155. This network was studied in detail considering the mRNA expression profiles in murine macrophages (miR-155 knock-out) responding to LPS [GSE77425; (Jablonski et al., 2016)]. Additionally, the role of miR-155 in macrophage polarization was also predicted by considering the miRNA-mRNA expression profiles in polarized and unpolarised murine macrophages (Lu et al., 2016). Subsequently, the coordinate biogenesis regulatory network of miR-125a-3p was also determined by considering mmu-miR-125a-3p as miRNA-1. Further, the probable role of miR-125a-3p as a CB-regulator was studied in macrophage polarization (Lu et al., 2016) and hematopoietic stem cell differentiation [GSE33691;(Gerrits et al., 2012)]. Moreover, miR-146a-5p co-ordinate biogenesis networks in murine monocytes and human monocytes in different LPS exposure conditions were determined utilizing miRNA expression profiling datasets [GSE87396, (Geng et al., 2016)][GSE125572,(Simmonds, 2019)] while has-miR-146a-5p CB-regulatory network in *Mycobacterium tuberculosis* infected macrophages was determined considering a previous study (Lin et al., 2017).

### Web-server for determining CB regulator-target relationships

The co-ordinate biogenesis data for each CB-regulator (miRNA) can be obtained from the COORD-BIO repository. Herein, the data mined during this work to study the co-ordinate biogenesis phenomenon is queried by considering a particular CB-regulator and expression profiles (miRNA and mRNA) in a scenario of interest. The front end of the server is html, PHP and java based while a perl script determines all possible CB regulator-target relationships for a particular CB regulator (miRNA) based on the conditions proposed for co-ordinate biogenesis phenomenon to occur. Further, context specific miRNA and mRNA expression profiling data may be utilized to determine the most likely sets of CB relationships. This web-server for determining probable CB miRNA pairs and associated secondary effector mRNA is freely available for use (http://www.hpppi.iicb.res.in/coordB2/index.html).

### Measurement of Nitric Oxide

Nitric oxide (NO) generation in response to LPS treatment post-anti-miR-146a transfection was quantified by Griess reaction as described previously (Ghose et al., 1999; Green et al., 1982). The amount of Nitric oxide produced by the cell was measure form culture supernatants after constructing standard curve with different concentrations of sodium nitrite. The concentration of Nitric oxide accumulated was expressed as μM NO.

### Cytokine measurement by ELISA

After completion of necessary experiments, culture supernatant has been collected. Using commercially available ELISA kits (R&D Systems), amount of TNF-α production under specific conditions were measure according to the manufacturer’s protocol.

### TUNEL assay

TUNEL assays were performed using DeadEnd™ Fluorometric TUNEL System kit (Promega) as per manufacturer’s protocol. Protocol has been described previously (Simmonds, 2019). Percentage of apoptotic cells has been counted analyzing multiple microscopic fields that has been represented on a bar graph.

## Acknowledgement

We acknowledge Witold Filipowicz and Susanta Roy Chowdhury for different plasmids constructs and also for valuable discussions. SNB and SC acknowledge CSIR-Indian Institute of Chemical Biology (IICB) for infrastructural support. We would like to thank Nucleome Informatics Private Limited for small RNA sequencing and analysis.

## Author contributions

SNB and SC designed the study, interpreted the data and prepared the draft manuscript. Bioinformatics data was generated and analysed by IM. S. Chatterjee and IM prepared the manuscript. All authors read and approved the final version of the manuscript.

## Data availability

miRNA sequencing data associated with this study is freely available in GEO. The data discussed in this publication have been deposited in NCBI’s Gene Expression Omnibus (Edgar *et al*., 2002; Barrett et al. 2013) and are accessible through GEO Series accession number GSE172473 (https://www.ncbi.nlm.nih.gov/geo/query/acc.cgi?acc=GSE172473).

